# Genetic, parental and lifestyle factors influence telomere length

**DOI:** 10.1101/2021.12.14.472541

**Authors:** Sergio Andreu-Sánchez, Geraldine Aubert, Aida Ripoll-Cladellas, Sandra Henkelman, Daria V. Zhernakova, Trishla Sinha, Alexander Kurilshikov, Maria Carmen Cenit, Marc Jan Bonder, Lifelines cohort study, Lude Franke, Cisca Wijmenga, Jingyuan Fu, Monique G.P. van der Wijst, Marta Melé, Peter Lansdorp, Alexandra Zhernakova

## Abstract

The average length of telomere repeats (TL) declines with age and is considered to be a marker of biological ageing. Here, we measured TL in six blood cell types from 1,046 individuals using the clinically validated Flow-FISH method. We identified remarkable cell-type-specific variations in TL. Host genetics, environmental, parental and intrinsic factors such as sex, parental age, and smoking are associated to variations in TL. By analysing the genome-wide methylation patterns, we identified that the association of maternal, but not paternal, age to TL is mediated by epigenetics. Coupling these measurements to single-cell RNA-sequencing data for 62 participants revealed differential gene expression in T-cells. Genes negatively associated with TL were enriched for pathways related to translation and nonsense-mediated decay. Altogether, this study addresses cell-type-specific differences in telomere biology and its relation to cell-type-specific gene expression and highlights how perinatal factors play a role in determining TL, on top of genetics and lifestyle.

## Introduction

With an increasingly ageing worldwide population, age-related diseases pose a great burden in clinical care and socioeconomics. Healthy ageing is the goal to counter this trend, but this term is complex and not easily defined. Several markers for premature or delayed ageing have been suggested, including telomere length^1, 2^, DNA methylation^3, 4^ and thymic function^5^. Telomeres are repetitive DNA structures located at the chromosome ends and, together with their associated proteins, play a fundamental role in chromosomal stability^6, 7^. Telomeres are known to decrease with age^2, 8^ as a result of multiple factors, including limiting telomerase activity^9^, the end-replication problem^1^, end-processing^10^ and oxidative stress^11^. Both genetic and environmental factors are known to influence telomere length. Several genetic loci have been associated with telomere length^12–16^, with heritability estimates ranging from 34%^17^ to 82%^18, 19^. However, heritability estimates often cannot distinguished between true genetic determinants and early life factors such as parental or environmental exposures that could affect telomere length during adulthood^18^. This is especially true in twin-based studies, where the early life exposures are confounded with genetic effects.

Importantly, most telomere length analyses carried out to date focused on blood leukocytes and did not explore cell-type- and tissue-specific variability. A recent study examining the variability of telomere length in a wide range of post-mortem tissues^20^ showed that, although whole blood telomere measurements might be a proxy for and synchronous with those of other tissues^21^, there are significant tissue-specific differences. However, this post-mortem tissue study did not address the possibility of telomere length differences between different cell types within a tissue. In addition, most large studies to date have used PCR or Luminex-based methods to measure relative telomere length in isolated genomic DNA, and these approaches have shown reproducibility biases^22, 23^ that could potentially explain the heterogeneous and contradicting findings^24^.

Telomere length may have important physiological consequences^25^. It has been proposed that telomere length might regulate gene expression^26–29^ but also that gene expression can directly contribute to telomere length attrition or conservation^30, 31^. Given the observed variability in telomere length with cell population^20^, it is conceivable that this will be related with cell-type-specific expression patterns, which have not been investigated to date.

Here, we explore telomere variation using Flow-FISH telomere length measurements in Lifelines Deep (LLD), a well-characterised population cohort from the Netherlands^32^. We measured telomere length in six different cell types in 1,046 participants. By combining this data with genetic information available for LLD participants and with rich phenotypic information that includes blood cell counts and immune markers, self-reported diseases, birth-related phenotypes, parental diseases and behaviour, epigenomics profiles and single-cell expression patterns in 62 individuals, we determined the major contributors to telomere length variation. Specifically, we studied: (1) the difference in telomere length across the six blood cell types, (2) the relationship between leukocyte telomere length with other ageing markers, (3) the correlation of telomere ageing markers with biochemical, parental and clinical phenotypes and mortality, (4) the contribution of genetic and non-genetic factors to variations in telomere length variation and (5) the cell-type-specific changes in gene expression associated with cell-type-specific telomere length variation, which may pinpoint major functional pathways related to telomere variability.

## Results

### Telomere length captures biological variability other than age

LLD^32^ is a population cohort from the northern Netherlands that includes participants with a wide age range (mean 43.9 years ± 13.7 sd, min 18.0, max 81.4) for whom we have deep phenotypic and molecular information [available data illustrated in **Supplementary Figure 1**, for descriptive statistics see **Supplementary Table 1**]. In 1,046 LLD participants, Flow-FISH^33^ was used to measure the telomere length of six blood cell types: granulocytes, lymphocytes, B-cells (CD45RA+CD20+), naïve T-cells (CD45RA+ CD20-), memory T-cells (CD45RA-) and NK-cells/fully differentiated T-cells (CD45RA+CD57+) (hereafter referred to as NK-cells) [**Figure 1A**]. We found that all six cell-type-specific telomere lengths decreased with age [**Figure 1B**] and were, on average, shorter in males than females over the entire age range [**Figure 1C**][**Supplementary Figure 2B** for all cell types]. We also observed a similar moderate negative correlation between age and telomere length among cell types (Pearson correlation, maximum r = -0.43, minimum r = -0.33) [**Figure 1D**]. These findings agree with those of a Flow-FISH‒based study in a North American cohort^9^, although the two cohorts differed in their participant recruitment selection criteria. Nevertheless, in the overlapping age ranges, both studies find comparable telomere lengths [**Supplementary Figure 2A**], which supports the accuracy of our measurements. We observe that naïve T-cells and B-cells have the longest telomeres on average, whereas NK- and memory T-cells show significantly shorter telomeres than other cell types (Paired Wilcoxon-test, p < 2×10^-16^. While naïve T-cells showed the highest mean telomere length [**Figure 1A**], they also showed the largest negative association of telomere length and age (linear model, slope of -0.034), from an average 8.59 Kb in the < 32.9 years age group (first quantile) to an average 7.32 Kb in individuals > 52.7 years (fourth quantile) [**Figure 1B**]. The rate of telomere loss we observe in naïve T-cells matches previous observations^9^ and does not support production of naïve T-cells from more primitive precursors after puberty and thymus involution^34^. Instead, we assume that naïve T-cells are maintained after puberty by homeostatic mechanisms that are likely to involve cell divisions that result in telomere loss, but these processes are currently poorly understood.

**Figure 1.**
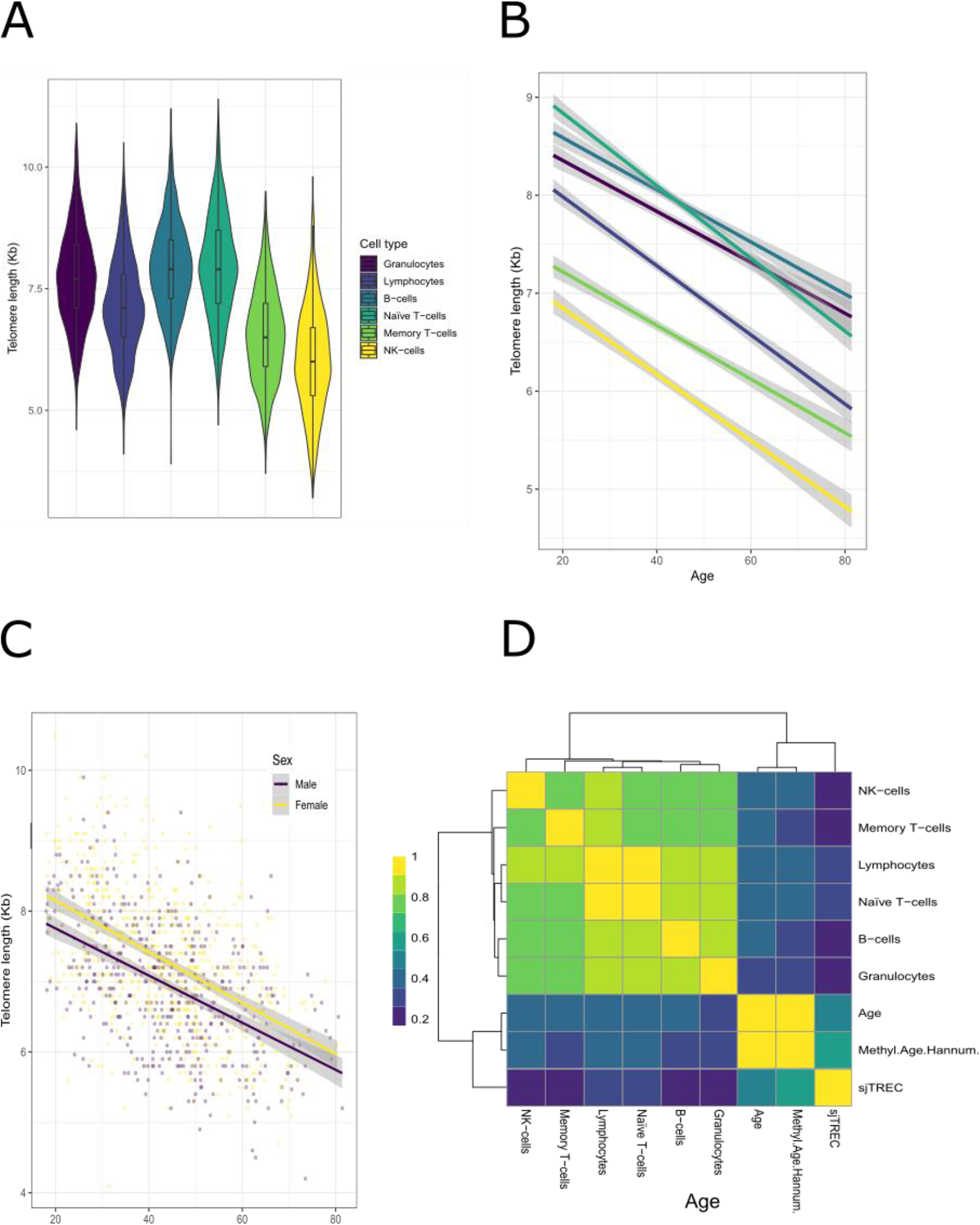
Telomere measurements in six blood cell types. **A.** Distribution of telomere length by cell type. **B.** Average trend line of telomere length. Grey shading indicates 95% confidence interval. Length decreases with age in all six cell types. Colour indicates cell type in both (A) and (B). **C.** Sexual dimorphism of telomere length (shown in lymphocytes). Trend line indicates average length per age. Grey shading indicates 95% confidence interval. **D.** Correlation (Spearman’s rho value) between the absolute value of telomere lengths, chronological age (Age), Hannum-based methylation age and sjTREC qPCR relative expression.

Next, we compared telomere length with other biological age markers, including the methylation-based Hannum^35^ age-index and signal joint T-cell receptor excision circles (sjTRECs) expression (CT values of a qPCR), which represents thymic TCR maturation for a given individual^36^ [**Figure 1D**]. Here, we observed that both methylation age and sjTREC were more strongly associated with chronological age than telomere length, and neither was highly correlated with telomere length [**Figure 1D**]. After removing the variability attributed to chronological age, methylation age and sjTREC were negatively associated (Pearson correlation, r=-0.36), but we found no association to telomere length and telomere lengths remained highly correlated between different cell types [**Supplementary Figure 2D**].

Overall, these findings suggest that telomere length captures biological variation other than chronological age, and the source of this variation is distinct compared to other ageing markers measured, specifically methylation age and thymic function.

### Genetic contribution to telomere length

To explore to what extent genetics can explain variation in telomere length, we first performed a heritability analysis. We used genotype data to infer genetic relations and fit a GREML model, while controlling for age and sex. This analysis provides an estimate of the total telomere length variability attributable to the additive effects of common genetic variants. The results show a median SNP-based heritability of 45%, with a maximum of 51% (naïve T-cells) and a minimum of 19.6% (NK-cells). Although the standard errors of the estimates were high, with a median of 22% [**Figure 2A**], the heritability estimate falls in a similar range to those of previous reports^37–39^ and the corresponding P-values were below 0.05 for all cell types except NK-cells (p = 0.15). This result supports the idea that genetic factors contribute significantly to telomere length and can partially explain the inter-individual variability. The higher genetic contribution to naïve T-cell telomere length that we observe might be explained by the fact that these cells reflect less environmental influence. On the other hand, environmental factors will impact telomeres in memory T-cells, including NK-cells, because antigen-mediated clonal expansion of these cells is typically triggered by environmental factors (e.g. infections), and thus telomere length will decrease due to the number of replications.

**Figure 2.**
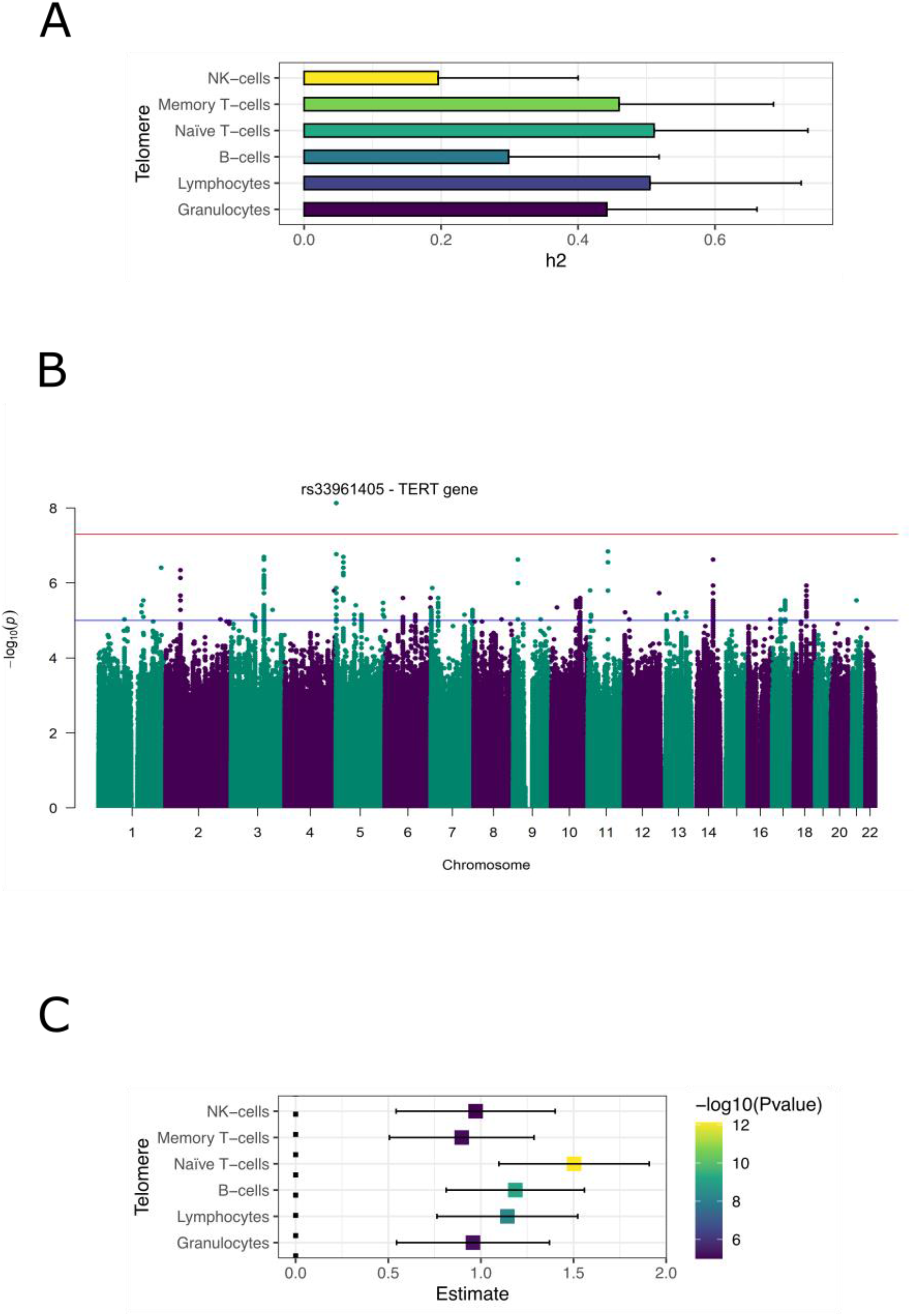
Genetic determinants of telomere length. **A.** Heritability estimation using a GREML model on the genotyped SNP data, from which a kinship matrix was estimated using GCTA^40^. The kinship matrix was used in a mixed model to estimate the variation explained by the genetic random effect. Units of heritability are represented as h2. **B.** Manhattan plot of genome-wide associations to telomere length from all cell types. Blue line indicates a suggestive significance threshold of 1×10^-5^. Red line indicates the standard genome-wide threshold of 5×10^-8^. The novel association with *TERT*, rs33961405 (5:1277577:G:A GRCh37), is labelled with its corresponding rsid. **C.** Correlation of the Polygenic Risk Score (PRS) of telomere length^15^ and telomere length of different cell types. Each square represents the estimated effect of the PRS in the cell type. Bars indicate 95% confidence interval.

To further explore the genetic contribution to telomere length, we performed a genome-wide association study (GWAS) of ∼7.5 million genotyped or imputed SNPs on telomere length in each cell type. First, we tested if we could replicate previously published loci associations. For this, we used the 20 genome-wide significant loci from a recent meta-analysis published by Li et al^15^. Of the 17 published SNPs for which we also had information, six were replicated in at least one cell type with a p-value < 0.05 and consistent allelic direction [**Supplementary Table 2.1**]. Similarly, we could replicate 6/12 European-associated genome-wide significant loci from a large trans-ethnic study^16^ [**Supplementary Table 2.1**]. We then looked at *de novo* associations and identified one significant association after applying permutation-based false discovery rate (FDR) correction: The A allele of rs33961405 (1000G EUR allele frequency = 0.49) located in an intron of the telomerase reverse transcriptase (*TERT*) gene was associated with decreased telomere length of T-cells (effect size = -0.19, SE = 0.03) [**Figure 2B**] [**Supplementary Figure 3**] (summary statistics of associations p < 1×10^-5^ presented in [**Supplementary Table 2.2**]). Other genetic variants located near the *TERT* gene were previously found to affect leukocyte telomere length; however, rs33961405 is novel and is in moderate linkage disequilibrium (LD) with a previously published lead SNP in *TERT* (rs2736100, LD r2 = 0.47 in 1000G EUR data)^12, 13^.

To reproduce previous GWAS associations on the individual SNP level, we computed a Polygenic Risk Score (PRS) from a large GWAS on telomere lengths^15^ (see Methods) and correlated it to our telomere length data. This analysis identified highly significant positive associations. The strongest genetic associations were seen in cell types with longer telomeres: naïve T-cells (linear model (lm), effect estimate = 1.5, p = 7.61×10^-13^) and B-cells (lm, effect estimate = 1.19, p = 6.78×10^-10^). Cell types with shorter telomeres, memory T-cells (lm, effect estimate = 0.89, p = 7.47×10^-6^) and NK-cells (lm, effect estimate = 0.97, p = 1×10^-5^), showed weaker associations of the genetic determinants. An exception to this trend was granulocytes, which showed a similar association range to B-cells and NK-cells (lm, effect estimate = 0.95, 6.02×10^-6^), despite having one of the longest average telomere lengths.

### Telomere ancestry ‒ parental age and smoking contribute to an individual’s telomere length

We exploited the extensive phenotypic information available for LLD study participants to uncover which environmental factors are correlated with telomere length. The phenotypic information consists of 90 different parameters, including blood parameters (e.g. leukocyte counts), anthropometric measurements (e.g. BMI), physiological parameters (e.g. blood pressure), various pre-existing diseases (e.g. hypertension or cancer) and lifestyle factors (e.g. smoking), as well as parental phenotypes and habits including parental diseases, smoking and age at participant’s birth [**Supplementary Table 3**].

To associate telomere length with different phenotypes, we built a linear model using telomere length as the dependent variable and the standardised phenotype measurement as the regressor, while controlling for age and sex. Of the non-genetic factors, blood cell counts were strongly associated with telomere lengths (**Supplementary Table 4.1**). Since cell counts might act as a confounder for other associations (as cell types have different telomere length and thus might confound the observed inter-individual differences), we included cell counts as covariates in the model. Using this new model, we identified 37 associations of 12 phenotypes with telomeres of any cell type using an FDR < 0.05 threshold (summary statistics can be found in **Supplementary Table 4.2**).

Several parental factors were consistently associated with telomere lengths in different cells. Smoking phenotypes such as ‘any parent smoking’, ‘father smoking during your childhood’ and ‘mother smoking during pregnancy’ were negatively associated with the telomere lengths of almost all cell types (with the exception of NK-cells). ‘Age of father when you were born’ and ‘age of mother when you were born’ were positively associated with the telomere lengths of four cell types. In a model combining both paternal and maternal age (on an l1 penalisation, see Methods), we found paternal age to shrink to near 0 while maternal age was kept constant (with the exception of granulocytes), highlighting that the effect of maternal age on telomere length is independent of paternal age.

Significant negative correlations with smoking were only observed with the father, both parents in combination and the mother during pregnancy. However, all cell types also showed nominally significant associations with maternal smoking. Maternal smoking associations showed weaker effect sizes than paternal (lm, 0.06 average difference, SE: 0.01, p = 3.2×10^-4^) and fewer participants had mothers who smoked than fathers who smoked (674 fathers vs. 381 mothers), factors which together may explain why maternal smoking did not reach FDR significance. Paternal and maternal smoking were, however, shown to have additive effects: A model that included a numeric variable describing the number of parents who were smokers showed stronger associations than binary smoking phenotypes (father, mother, or any parent).

We further analysed the associations to other available smoking phenotypes. Here we found a consistent nominally significant negative effect of ‘father smoking’, ‘mother smoking’ and ‘mother smoking during pregnancy’ in all the cell types tested. In addition, passive smoking during an individual’s lifetime also influenced telomere length, with the factor “do people smoke near you at work” associated to shorter telomeres in 5/6 cell types. Conversely, current smoking of the participant did not reach nominal significance in any cell type [**Supplementary Figure 4A**].

In addition, we found four negative associations with participant BMI and three with participant waist circumference. These negative associations with BMI and waist circumference could be driven by their correlations to other phenotypes. We therefore explored the association of BMI with other phenotypes in a larger cohort of 10,000 participants from the same population^41^ and observed a positive correlation between BMI and parental smoking (Pearson, p < 4.78×10^-6^) and a negative correlation between BMI and parental age (Pearson, p < 9.15×10^-7^). After accounting for potential confounding effects of parental age and smoking habits, the associations between telomere length and BMI phenotypes remained significant [**Supplementary Figure 4B**]. This observation supports the conclusion that the associations of telomere length to BMI are not driven by the confounding effects of parental age or smoking.

In addition, we found one cell-specific (having one cell type below FDR < 0.05) positive association with poorly healing wounds (granulocytes) and three cell-specific negative associations with blood alpha-1 antitrypsin (AAT) (Memory T-cell), pulse rate (granulocytes) and weight (B-cells) [**Figure 2A**] (summary statistics in **Supplementary Table 4.2**).

Finally, we assessed how much of the variation in telomere length not attributable to participant age could be explained by intrinsic, parental and genetic factors. For environmental factors, we included all associated factors with an FDR < 0.05 in at least two cell types. Genetic factors were addressed using PRS from a recently published telomere length GWAS^15^, see Methods. We fitted four nested models and estimated the added variability (R2) explained by each (see Methods) [Figure 3B]. This revealed that most of the variability is explained by the addition of sex, BMI, waist circumference and cell counts (from 3.2 to 8%), depending on the cell type. The addition of parental phenotypes (parental age and smoking) added less information (0.8%), on average, than the multiple intrinsic factors. Finally, the contribution of genetics (average 3.7%) was lower than that of intrinsic factors (average 5.9%, considering sex as an intrinsic factor rather than a genetic one) in most cells, which is in line with the larger impact of environmental effects that we observed in the heritability analysis.

**Figure 3.**
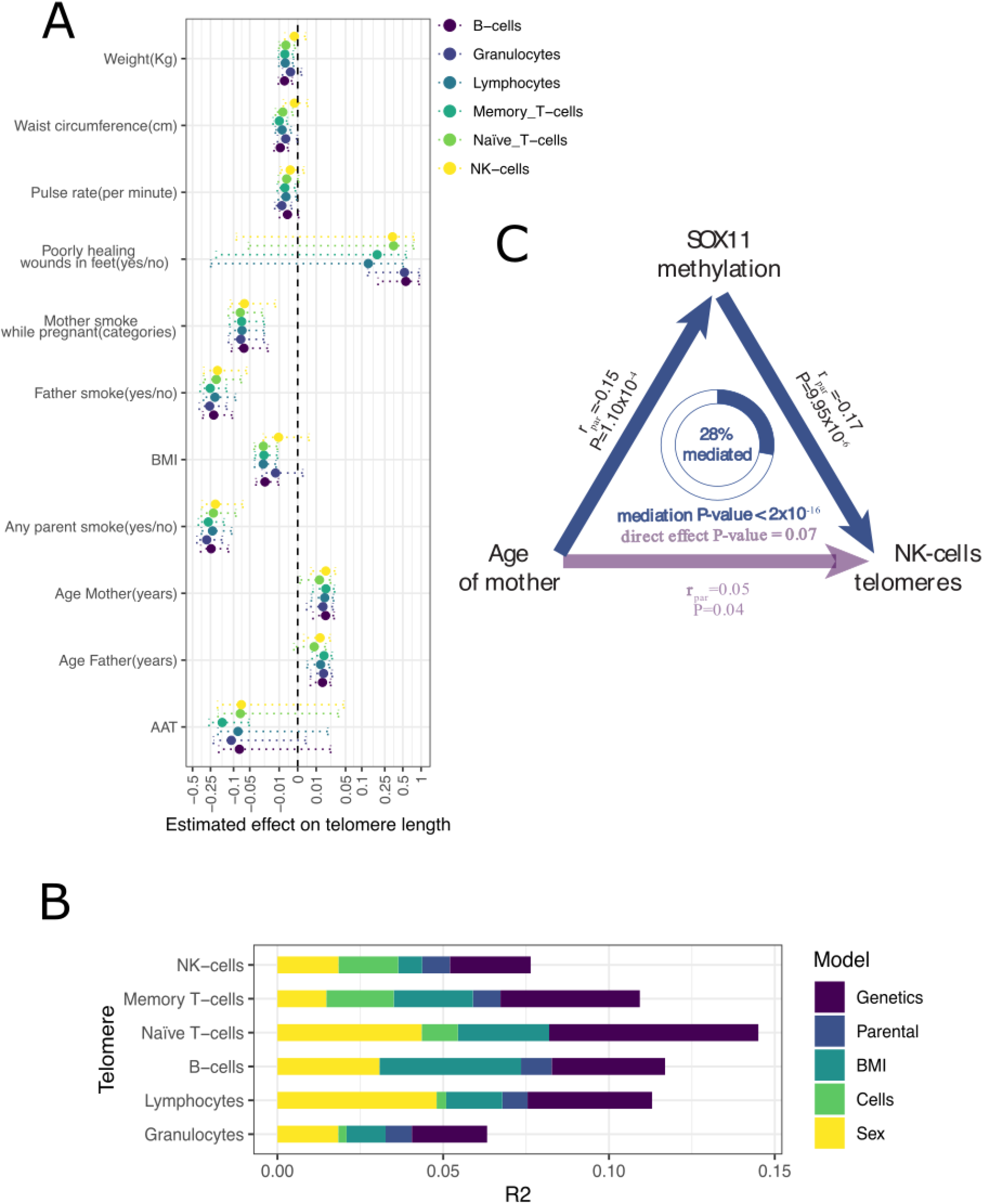
Non-genetic factors contribute to telomere variation. **A.** Phenotype effect on telomere lengths of different cell types (with at least one significant association). Dashed lines show 95% confidence interval (estimate ± 2xSE). X-scale is symmetrical log-transformed (denominator constant = -2). BMI: Body mass index, AAT: Alpha-1 Antitrypsin Test. **B.** Total variance of the telomere length explained after removing the effect of age. Only phenotypes associated with at least two cell types (FDR < 0.05) are used. Colour indicates the different partitions of variability. **C.** Mediation effect of methylation of *SOX11* in the maternal age effect on NK-cell telomere length variability.

### Epigenetic changes may mediate the effect of parental phenotypes on telomere length

It has been proposed that maternal and paternal phenotypes, such as smoking habits and age at pregnancy, may affect a child’s phenotypes by inducing changes in methylation levels^42–47^. Several studies have reported an effect of parental age and smoking on the “methylation age” of the child^43, 44, 46, 48^. We therefore investigated whether the effect of parental phenotypes on telomere length could be mediated by changes in methylation levels. We first performed a GWAS of parental smoking and age with DNA methylation levels available for the same samples (N = 749). This resulted in 19 genome-wide significant associations of methylation probes with parental phenotypes at a Bonferroni-corrected p < 0.05 (summary statistics in **Supplementary Table 5**). Combining these results with the associations of parental phenotypes with telomere length (p < 0.05) and of telomere length with methylation (p < 0.05), we performed mediation analysis for the 17 resulting triplet associations. This analysis identified four triplets that showed significant mediation of the parental phenotype effect on telomere length through methylation of various probes (see Methods) [**Supplementary Table 5**]. All these significant mediation results were for the age of mother, even though we tested fewer triplets for maternal (6 triplets) than paternal age (11 triplets). Maternal age effects thus appear more likely to be mediated by methylation than paternal age effects.

One interesting example of this methylation mediation is for the SRY-related HMG-box gene *SOX11*. This gene is a known transcription factor and proliferation gene that plays an important role in embryonic development, cell fate determination and cancer. Our results indicate that the positive association of maternal age at birth with telomere length in NK-cells may be mediated by decreased methylation of CpG islands located in the promoter of *SOX11* (mediation p < 2×10^-16^), with up to 28% of the effect mediated through methylation of *SOX11* [**Figure 3C**] [**Supplementary Table 5**].

### Telomere length changes show cell-type-specific associations with gene expression level

To study the relationships between telomere length variation with gene expression changes, we used single-cell RNA-sequencing (scRNA-seq) data generated on cryopreserved peripheral blood mononuclear cells (PBMCs) from 62 LLD donors, for which telomere length on six cell types was measured^49^. To classify cells, we combined the high resolution cell-type-annotations by Azimuth^50^ to closely reflect the resolution of the Flow-FISH annotations (i.e. naïve and memory CD4T and CD8T cells, NK- and B-cells) (see Methods) (**Supplementary Table 6**). First, we confirmed that the subset of 62 LLD donors had similar telomere length distributions to the entire study population (1,046 LLD donors) (**Supplementary Figure 5**). We then performed telomere length differential gene expression analyses at single-cell resolution (sc-DEA) by selecting the matched telomere length measurement and gene expression level for each of these cell types (see Methods) (**Supplementary Table 6**). These analyses revealed *DNAJA1* to be positively associated with telomere length in memory CD8T cells (effect size = 0.05 log-fold change (LFC) per telomere length unit, FDR = 0.03) (**Supplementary Table 7.1**). *DNAJA1* encodes a heat shock protein 70 co-chaperone that was previously reported to bind telomeres in a study that used *in vivo* cross-linking, tandem affinity purification and label-free quantitative mass spectrometry^51^.

Because we only identified one differentially expressed (DE) gene, we wondered whether this could be due to insufficient statistical power. To address this, we performed differential gene expression analysis on T-cells combining multiple cell types together (Methods) while controlling for cell type annotation (**Supplementary Table 6**). This strategy increased the number of cells per donor and thus statistical power. However, in contrast to analysing each cell type separately, genes identified with this combined strategy will likely have a similar association with telomere length across cell types. We identified 97 unique DE genes, one in CD8T cells, 44 in CD4T cells and 91 in all T-cells (**Figure 4**) (**Supplementary Table 7.2**, **Supplementary Figure 6**), including the *DNAJA1* association reported in our previous analysis

**Figure 4.**
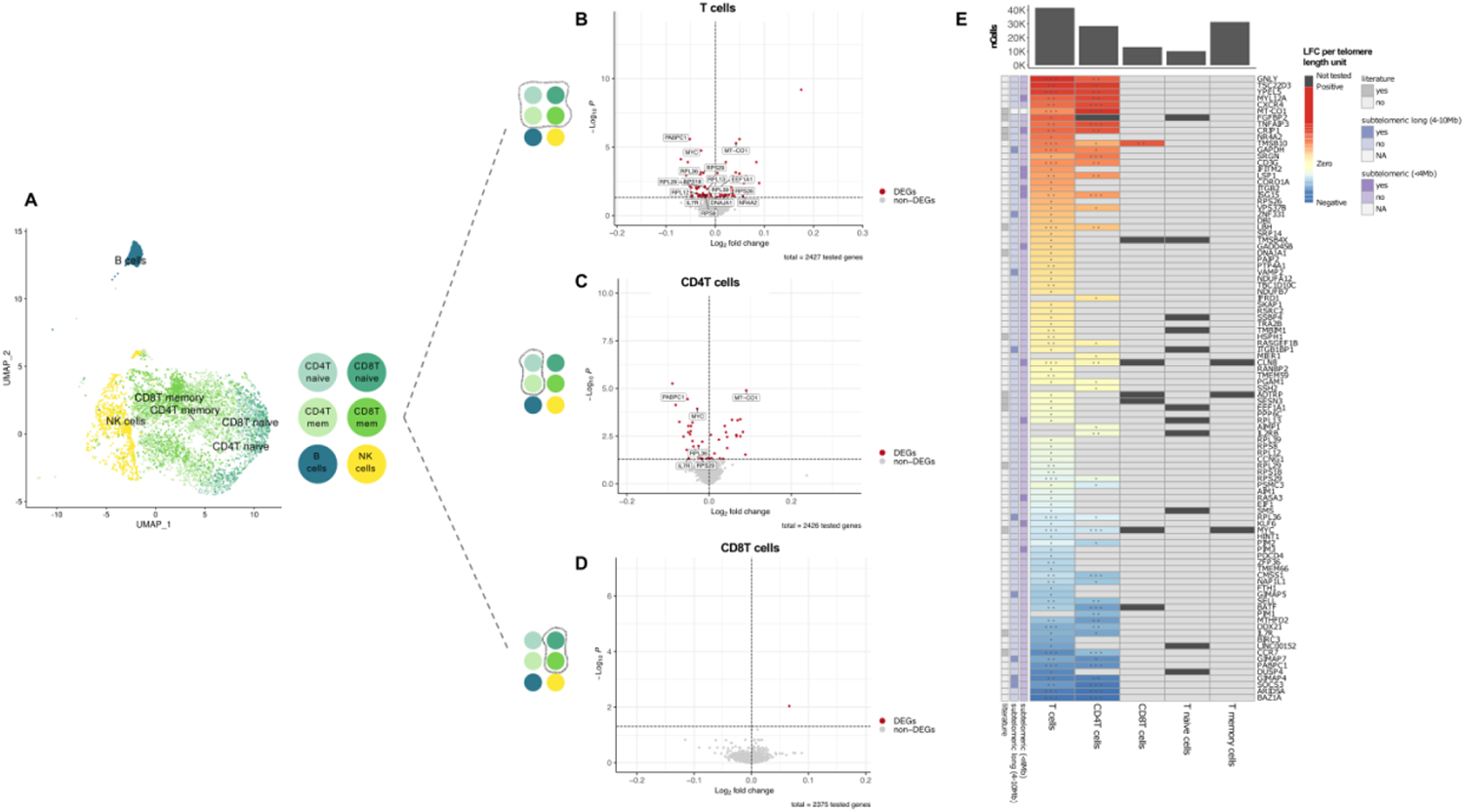
Differential gene expression changes with telomere length across cell types. **A.** UMAP plot of the cells from the subset of 62 LLD donors with both scRNA-seq and Flow-FISH telomere length data. The cells are coloured by the cell type classification that closely reflects the resolution of the Flow-FISH annotations (i.e. naïve and memory -mem- CD4T and CD8T cells, NK- and B-cells). For visualisation purposes, we down-sampled each of Azimuth’s predicted cell types to 500 cells. **B, C and D.** Volcano plots showing the results of DE approach II in T-, CD4T and CD8T cells, respectively. For DE approach II, we combined multiple cell types together (in A) in the same analysis (i.e. combining all T, all CD4T, all CD8T, all naïve T and all memory T-cells). DEGs with telomere length are represented in red. Non-DEGs are represented in grey. Labels correspond to the DEGs mentioned in the text. **E.** Heatmap of log-fold change (LFC) per telomere length unit for the set of 97 unique DEGs identified in T-, CD4T and CD8T cells. Non-DEGs are shown in light grey. Genes not tested (i.e. those below 10% expression cut-off) are shown in dark grey. The significance level of the DEGs corresponds to the following arbitrary FDR thresholds: FDR<0.001 (***), FDR<0.01 (**) and FDR<0.05 (*). The genes are sorted by their average LFC across cell types. The row annotation bars show whether the genes are located at the subtelomeric (<4Mb) or subtelomeric long (4–10Mb) region and if the genes were previously reported in any of the following studies: Pellegrino-Coppola et al., 2021^53^, Tacutu R et al., 2018^54^, Buxton JL et al., 2014^29^ or Nittis T et al., 2010^51^ (**Supplementary Table 8**). The distance to the telomeres was not calculated for the mitochondrial gene (MT-CO1) (subtelomeric (<4Mb) and subtelomeric long (4–10Mb) = NA). The column annotation on the bar plot shows the total number of cells (nCells) per cell type.

**Figure 5.**
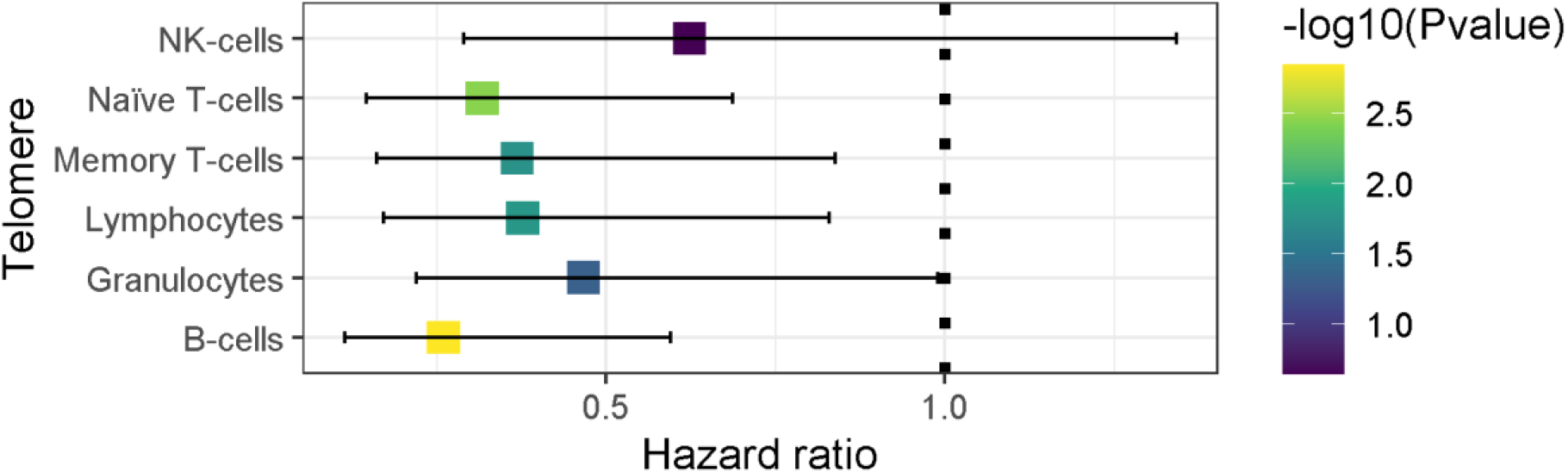
All-cause mortality risk of telomere length. Y-axis represents the hazard ratio of all-cause mortality estimated by a Cox regression on each cell type’s telomere length. Bars indicate estimated 95% confidence interval.

Next, we explored potential mechanisms explaining the telomere length associations with expression in the set of 97 genes. Several mechanisms have been described by which telomere length may affect gene expression levels. The first is the telomere position effect (TPE)^26^. This mechanism results in decreased repression of genes located in the subtelomeric region when telomeres shorten^27^ and in subsequent overexpression, which is a process associated with CpG-methylation^29^. To determine whether our sets of DE genes were influenced by TPE, we tested whether there was an enrichment of DE genes in the subtelomeric region (< 4Mb from chromosome end) (**Supplementary Table 8**). In CD4T cells, we found a significant enrichment of genes positively (5/19, p = 0.007) but not negatively associated (0/24, p = 1) associated with telomere length (**Supplementary Figure 7**). We did not observe any enrichment in all T-cells (8/45, p = 0.05 for positively associated and 4/45, p = 0.58 for negatively associated genes) (**Supplementary Table 8**). The enrichment we observe in the CD4T cells is opposite to what we expected, i.e. we expected to identify genes that were negatively associated with telomere length as a result of the lost repression due to shortening. Our contrasting findings suggest that other mechanisms might be at work in the associated genes.

A second mechanism by which telomere length may affect gene expression levels occurs through a telomere position effect over long distances (TPE-OLD). Such long-distance gene expression regulation was shown to be the result of telomeres forming chromatin loops with enhancer and repressor regions of genes within 10Mb of chromosome ends^28^. These interactions are dependent on telomere length but are not linearly related with shortening of the telomeres. We did not find any enrichment within our DE gene sets for genes acting through a potential TPE-OLD mechanism (i.e. 4–10Mb from chromosome end) in either T-cells (4/45 DE genes positively associated with telomere length, p = 1; 5/45, DE genes negatively associated with telomere length, p = 0.8) or CD4 cells (1/19 DE genes positively associated with telomere length; p = 0.7; 4/24 DE genes negatively associated with telomere length, p = 0.52) (**Supplementary Figure 7**) (**Supplementary Table 8**). However, the generalised linear mixed model (GLMM) we used for sc-DEA assumes linearity and therefore might not allow us to properly test this hypothesis, which assumes non-linear relations.

Since many of the identified DE genes did not fall into the TPE or TPE-OLD categories, other unknown effects on gene expression through telomere length may be at play or, conversely, the expression of specific genes may affect telomere length. For several DE genes, we found additional evidence for a telomere connection in literature (**Figure 4**) (**Supplementary Table 8**). Firstly, three of our hits (*DNAJA1*, *EEF1A1* and *RPL29*) were previously reported in a screen for novel telomere binding proteins, indicating their direct involvement in telomere maintenance^51^. Secondly, CpG methylation of *NR4A2* was previously found to be associated with telomere length in whole blood^29^. *NR4A2* is involved in T-cell maintenance through regulation of Treg suppressor functions and repression of aberrant Th1 induction^52^. The third line of gene–telomere interaction evidence is the overlap with leukocyte ageing– associated genes that were previously identified in whole blood after a deep correction for cell type composition (8 out of 84 testable genes overlapped; odds ratio = 3.3, p = 6×10^-3^)^53^. Finally, four genes also overlapped with the GenAge human database, the core Human Ageing Genomic Resources (HAGR) database composed of >300 human ageing–related genes^54^. These four were related to immunosenescence^55, 56^ (*IL7R*), cellular senescence^57, 58^ (*MYC*) and longevity ^59–62^ (*EEF1A1* and *MT-CO1*), in line with telomere shortening being a well-known hallmark of both cellular senescence and organismal aging^63, 64^. However, we did not find a significant overlap between the genes we identified and the ones reported in that study (odds ratio = 1.4, p-value = 0.53).

After exploring individual gene associations, we wondered if the identified DE genes belonged to similar functional pathways and could thus highlight the biological interplay between telomeres and gene expression. To explore this, we performed a functional enrichment analysis within the different sets of DE genes. In CD4T cells, the JAK-STAT signalling pathway was negatively associated with telomere length (enrichment ratio = 18.63, FDR = 4×10^-3^) (**Supplementary Table 9**). This pathway has previously been associated with telomerase regulation in haematologic malignancies^65^ and immunosenescence^56^. Among the DE genes involved in the JAK-STAT signalling pathway, interleukin-7 receptor (IL7R) plays a critical role in lymphoid cell development^66, 67^, and its gene expression network has been proposed as a potential biomarker for healthy aging^55^. In all T-cells, we identified 26 pathways enriched among the genes negatively associated with telomere length, many of them related to translation, including peptide chain elongation, eukaryotic translation elongation or termination and initiation, among others (**Supplementary Table 9**). Previous genetic screens in yeast^68–71^ and *Arabidopsis thaliana*^72^ identified ribosome biogenesis as one of the largest gene categories linked to telomere length. On top of that, in human fibroblasts, cellular senescence triggered through telomere shortening can diminish ribosome biogenesis, resulting in rRNA precursors and accumulation of ribosomal proteins (such as RPL29, which we found to decrease expression with telomere length in T-cells)^73, 74^. In addition, our set of negatively enriched pathways in T-cells also revealed the nonsense-mediated decay (NMD) pathway (including *PAMPC1* expression^75^) that has been recently proposed to regulate the levels of specific mRNAs that are important for telomere functions^76^.

### Short telomere length associates with all-cause mortality, independent of age

The most apparent implication of having abnormal telomere length may be its effect on morbidity and mortality. We collected all-cause mortality data for the previous 8 years, i.e. since the start of LLD data collection. Eleven study participants had died at the time of the analysis, and we used a Cox model while controlling for cell counts, age and sex to assess the predictive power of telomere length (using the survival time measured as days as an outcome). Despite the very low sample size, we found a consistent association of shorter telomeres and higher all-mortality death risk. The telomere length from all cell types except NK-cells reached statistical significance [**Figure 4**][**Supplementary Table 10**], indicating the effect of the telomere length is independent of the effect of age in all-death prediction.

## Discussion

This work presents the largest study to date on telomere length variation in a population cohort based on cross-sectional Flow-FISH^33^ measurements. We measured telomere lengths in six immune-related cell types from 1,046 participants from the northern Netherlands LLD cohort^32^. We investigated the effect of genetics and 90 phenotypes, including parental factors and a wide range of general and environment factors, on telomere length. This identified the most important factors that influence telomere length and, based on our findings, we propose potential mechanisms of action for some of these factors. Finally, we identified cell-type-specific transcriptional modules related with immuno-senescence that might explain the downstream effects of telomere length attrition.

Previous studies of telomere length variation in population cohorts have mainly relied on classical terminal restriction fragment measurements^77^, qPCR-based methods^78, 79^ or computational methods^16^ to measure telomere length. In the current study, we used Flow-FISH in a large epidemiological setting. Flow-FISH allows accurate measurements of the average telomere length in different cell types^80^. This allowed us to observe the distribution of telomere length in different cell types in addition to the more commonly studied telomere length of total leukocyte population^15, 18, 19^. Our observations reveal a similar trend of telomere length decrease with age and a high correlation in telomere length between different cell types (Pearson correlation r minimum 0.70, mean 0.81, maximum 0.94). These finding overlap with previously published between-tissue telomere length correlation ranges (leukocyte, skin, skeletal muscle and subcutaneous fat, Pearson r from 0.76 to 0.88)^81^ but show a smaller range than those reported in post-mortem tissues (highest Pearson r = 0.40 between transverse and sigmoid colon)^20^. Nevertheless, it must be noted that measurements were carried out using Southern blot in^81^, whereas the measurements in^20^ where Luminex-based. These findings agree with previous studies that indicated that despite having different telomere lengths in adults, the overall effect of age on telomere length is similar in different cell types^81^. Additionally, our results show high reproducibility with respect to a previous Flow-FISH population study in a North American cohort^9^.

The average telomere length in each cell type is a complex phenotype that reflects cell-type-specific, genetic, intrinsic and environmental effects. Many previous studies have tried to estimate the role of genetics in explaining the variation of telomere length in humans^18^, producing a wide range of estimates. These differences suggest there are potential environmental effects confounding the heritability estimates. For instance, sibling-based studies^17, 38, 82^ have the disadvantage of not being able to distinguish between genetics and early life or prenatal exposures. In our study, which more closely resembles approaches that estimate “chip-based” heritability in unrelated individuals^83, 84^, we estimated narrow sense heritability in six different cell types based on a mixed-model approach using sample-to-sample genotype kinship estimations (GREML)^40^. One advantage of this study is that LLD participants were specifically selected to have a common ancestry but not be highly related among each other, meaning that the GREML estimation should not be biased by those factors. After accounting for age and sex, our GREML estimation indicates a median heritability of 40%, which is in the same range as previously reported^85^. However, our estimate does not include the X or Y chromosomes, which might play a role in telomere heritability^86^. In addition, the relatively small sample size of our study (∼1,000) hampers the accuracy of our estimation, and we observe large standard error intervals. Nevertheless, the heritability levels that we find are indicative of a large amount of telomere variability being caused by environmental factors, which contrasts with other studies that identified extremely high telomere heritability (0.7^19^, 0.78^82^, 0.81^87^). In our GWAS analysis, we identified one intronic SNP in the TERT gene that was associated at 5×10^-8^ and in moderate LD (0.47) with previously reported associations in this locus. The GWAS analysis in our cohort was limited by our relatively small sample size (911 individuals), but PRS scores from a recent large telomere GWAS^15^ indicate strong association of previously identified genetic variants with telomere length of all cell types.

Despite the on-going efforts to understand the effect of genetics on telomere length^12–16^, a large fraction of unexplained variability remains, which may be attributed to environment. We therefore investigated the relation of 90 phenotypes with telomere length. This analysis pointed to a role for parental phenotypes in the telomere variations in their children. Higher paternal age has previously been associated with longer telomere lengths in humans^19^. Our results agree with this but also identify a maternal age effect on telomere length. Given the high correlation between paternal and maternal age (r = 0.77), the maternal age effect we observe might be confounded by paternal age. It is known that sperm telomere length increases with age, whereas telomere length in somatic tissues, including leukocytes, diminishes with age^88, 89^ providing a potential explanation for longer leukocyte telomere length in offspring of older fathers. This effect might even be additive over consecutive generations^90^. In addition to parental age, we identified a novel telomere association with parental habits in which parental smoking was negatively associated with telomere length. This effect could either be prenatal or early life.

Additionally, even with the possibility of being caused by common genetic architecture, parental effects in offspring are more plausibly caused by a mechanism other than genetics, such as epigenetic modifications. We therefore tested whether methylation might mediate the effect of parental phenotypes such as age and smoking on telomere length in their children. The effect of maternal age, and not paternal age, on telomere length was mediated through methylation of several genes, suggesting that the maternal age effect is not completely confounded by paternal age. Here we highlighted the mediation of telomere length in NK-cells via the methylation of *SOX11*. It is not clear how *SOX11* methylation may lead to telomere shortening, as *SOX11* is mostly expressed during embryonic development. Previous studies have shown that promoter hypermethylation of *SOX11* inhibits *SOX11* expression in cancer cells^91^. In addition, several studies have shown an anti-senescent effect of *SOX4*, which belongs to the same *SOXC* group as *SOX11*^92^. It is important to stress, however, that this is a proposed mechanism, and that experimental validation will be needed to resolve the nature of such associations.

Apart from parental factors, we identified that women have longer telomeres than men, as previously identified^86, 93^. In addition, BMI was associated with shorter telomeres^20, 94^. However, we did not observe other previously reported associations, such as with participant smoking habits^20, 95^. Combining all identified intrinsic, parental and genetic factors other than age with significant effects at least in two cell types, we estimate that only 10%, on average, of the observed telomere length variability can be explained, with only about 3.7% attributable to genetics (without accounting for the genetics factors that cause sex differences). These findings show that environmental and intrinsic effects have a greater impact than genetics on telomere length variation. Nevertheless, it is worth noting that the model applied in this study does not consider interactions between the different layers of information, where genetics could potentially impact the effects of the other environmental features. In addition, we are only considering highly associated additive SNPs and no other possible epistatic relationships, which means we are probably only setting a lower bound on the amount of variability that we can account for.

Our single-cell transcriptomic analyses identified a set of 97 unique genes that are significantly associated with telomere length across T-cell types. Three of them (*DNAJA1, EEF1A1* and *RPL29*) were previously revealed as telomere binding proteins^51^, indicating that our approach captures genes directly involved in telomere length dynamics. Moreover, our study provides additional insight into the direction of effect. For example, we found *RPL29* to be negatively associated with telomere length, which is in line with previous studies describing accumulation of RPL29 as a biomarker of senescence^73^. Looking at the broader context, we observed functional enrichment of genes involved in translation and the NMD pathway within the gene set negatively associated with telomere length in T-cells, which might have important physiological consequences.

Even though some of the telomere length–associated genes were located near the telomere ends (i.e. 12 and 9 DE genes that fall < 4Mb and between 4–10Mb from the telomere ends, respectively), in general, our differential expression findings could not be explained by previously described mechanisms (TPE^26, 27^ and TPE-OLD^28^). This suggests that there are other mechanisms by which telomere length regulates gene expression, or vice versa^51^. Follow-up experimental perturbation studies will be required to make this distinction, for example by inducing telomere shortening (replicative senescence) followed by gene expression analysis or by inducing gene knockdowns followed by telomere length measurements.

Finally, by making use of longitudinal data on participant survival, we were able to replicate a negative association between telomere length and all-cause mortality^96^, although the small number of deceased participants (11) hampers the wider interpretation of these results. Further studies of Lifelines participants will clarify this predicted relationship^97^.

## Conclusions

Telomere length is a complex trait in human populations whose variability is influenced by genetics, lifestyle, intrinsic and environmental factors. Identification of such effects has been hampered by low reproducibility in measurements, limited phenotypic information about study participants and lack of cellular resolution to study cell-type‒specific nuances. Applying reproducible methodology (Flow-FISH), we show that there is substantial variation in telomere length among blood cell types that are usually measured together. We find a consistently lower telomere length in males and determine the influence of genetics, intrinsic phenotypes and parental phenotypes and habits. In particular, our identification of a novel effect of parental smoking habits highlights the importance of accounting for common environmental influences usually overlooked in family-based telomere length heritability estimates. Additionally, we propose a possible epigenetic mediation of maternal age effects on offspring telomere length. We also linked telomere length variability with transcriptomic differences at the single-cell level. These results suggest that T-cells are sensitive to telomere length–induced expression changes that can act through both short- and longer-range interactions, but also that cell expression may directly influence telomere dynamics. Through gene expression regulation, telomere shortening may contribute to ageing and age-associated diseases long before critical DNA damage starts. Finally, we confirm telomere length as an independent risk factor for death, which emphasises the importance of understanding the source of its variability in order to develop healthier life habits.

## Acknowledgments

We thank J. Dekens for management, A. Maatman and M. Platteel for technical support and K. Mc Intyre for English editing. This project was funded by the BBMRI grant for measuring telomere length and by a Rosalind Franklin Fellowship to A. Zhernakova. The researchers participated in this project are supported by Netherlands Heart Foundation (IN-CONTROL CVON grants 2012-03 and 2018-27); the Netherlands Organization for Scientific Research (NWO) Gravitation Netherlands Organ-on-Chip Initiative to J.F. and C.W.; NWO Gravitation Exposome-NL (024.004.017) to J.F., A.K. and A.Z.; NWO-VIDI (864.13.013) and NWO-VICI (VI.C.202.022) to J.F.; NWO-VIDI (016.178.056) to A.Z.; NWO-VIDI (917.14.374) to L.F.; NWO-VENI (194.006) to D.V.Z.; NWO-VENI (192.029) to M.W.; NWO Spinoza Prize SPI 92-266 to C.W.; the European Research Council (ERC) (FP7/2007-2013/ERC Advanced Grant 2012-322698) to C.W.; ERC Starting grant 637640 to L.F.; ERC Starting Grant 715772 to A.Z.; ERC Consolidator Grant (grant agreement No. 101001678) to J.F.; and RuG Investment Agenda Grant Personalized Health to C.W. MM work is supported by RYC-2017-22249 and PID2019-107937GA-I00 grants. T.S. holds scholarships from the Junior Scientific Masterclass, University of Groningen[grant no. 17-34]. AR is funded by a Formación Personal Investigador (FPI) grant [grant no. PRE2019-090193].

The Lifelines Biobank initiative has been made possible by a subsidy from the Dutch Ministry of Health, Welfare and Sport; the Dutch Ministry of Economic Affairs; the University Medical Center Groningen (UMCG, the Netherlands); the University of Groningen and the Northern Provinces of the Netherlands. The authors wish to acknowledge the services of the Lifelines Cohort Study, the contributing research centres delivering data to Lifelines and all the study participants.

## Material & Methods

### 1. Lifelines Deep cohort

#### Phenotype data description

Lifelines is a multi-disciplinary prospective population-based cohort study examining, in a unique three-generation design, the health and health-related behaviours of 167,729 persons living in the North of the Netherlands. It employs a broad range of investigative procedures to assess the biomedical, socio-demographic, behavioural, physical and psychological factors that contribute to the health and disease of the general population, with a special focus on multi-morbidity and complex genetics^41^. We collected data from the subcohort Lifelines DEEP (LLD, n = 1,057, 57.6% female, mean age (including months) 43.9 years [range 18–81.4 years])^32^. Extensive information on demographics, health and lifestyle factors including smoking and diet was collected via detailed questionnaires. Mean BMI of participants was 25.1 [range 15.8‒44.9]. Common age-related diseases within the cohort included hypertension (23% of 841 participants with information), type 2 diabetes (1.3% of 1,039 participants with information) and hypercholesterolemia (14% of 900 participants with information). In the cohort, 20% of individuals smoked currently, 48% smoked for at least 1 year and 37% had mothers and 65% had fathers who smoked.

#### Omics data description

Genome-wide genotyping data was generated as described in^32^. Genotype data processing was done as described in^98^. Briefly, microarray data were generated on CytoSNP and ImmunoSNP and then processed on the Michigan Imputation Server^99^ to perform phasing using SHAPEIT and imputation using HRC version R1 as reference^100^. We excluded SNPs with imputation quality r^2^ < 0.5, minor allele frequency (MAF) < 0.05, call rate < 0.95 and Hardy-Weinberg equilibrium test P < 1×10^-6^, which resulted in 5,327,634 SNPs used in subsequent analyses. Genotype data was available for 911 samples with non-missing telomere measurements.

Genome-wide methylation data generation for this cohort was described previously^101^. Briefly, the EZ DNA Methylation kit (Zymo Research) was used to bisulfite-modify 500ng of genomic DNA, which was hybridised on Illumina 450K array. Methylation probes were remapped to ensure their correct genomic location, and probes with known SNPs at the single base extension site or CpG site were removed. Next, data was processed using a pipeline described by Tost and Touleimat^102^. We used DASEN-normalised data with subsequent quantile normalisation and probe scaling applied. Methylation data for 418,499 probes was available for 651 samples with telomere measurements in at least one cell type.

### 2. Flow-FISH Telomere length measurement

Telomere length measurements using automated multicolour flow-fluorescence in situ hybridisation (Flow-FISH) was performed as described^33^. Briefly, white blood cells were isolated by osmotic lysis of erythrocytes in whole blood using NH4Cl. White blood cells were then mixed with bovine thymocytes of known telomere length (which served as an internal control), denatured in formamide at 87°C, hybridised with a fluorescein-conjugated (CCCTAA)3 peptide nucleic acid (PNA) probe specific for telomere repeats and counterstained with LDS751 DNA dye. The fluorescence intensity in granulocytes, total lymphocytes and lymphocyte subsets defined by labelled antibodies specific for CD20, CD45RA and CD57 relative to internal control cells and unstained controls was measured by flow cytometry to calculate the median telomere length from duplicate measurements.

Out of a total 1,388 participants, we could not measure the telomere length in any cell type in 207. In addition, granulocytes could not be measured in 109 participants, B-cells could not be measured in nine participants and NK-cells could not be measured in 17 participants (all of these were different participants). We decided to remove all participants with at least one missing cell type, which resulted in a final subset of 1,046 participants.

### 3. sjTREC measurement

DNA from whole blood was isolated using the conventional protocol with Proteinase K digestion followed by phenol extraction and isopropanol precipitation. Next, we performed a TaqMan quantitative real-time PCR approach to quantify sjTREC expression (signal joint excision circles produced during T-cell development) using ViiA™ 7 Real-Time PCR System (Life technologies). TaqMan qPCR was performed on 75-100 ng DNA in a 12 μl reaction mixture containing 18 µM of each primer for sjTREC (5’-TCGTGAGAACGGTGAATGAAG-3’ and 5’-CCATGCTGACACCTCTGGTT-3’) and for albumin as housekeeping gene (5’-TGAACAGGCGACCATGCTT-3’ and 5’-CTCTCCTTCTCAGAAAGTGTGCATAT-3’) and 5 µM of hydrolysis sjTREC probe 5’-(FAM) CACGGTGATGCATAGGCACCTGC-3’ (TAMRA) and albumin probe 5’-(FAM) TGCTGAAACATTCACCTTCCATGCAGA -3’ (TAMRA). PCR runs started with incubation at 50°C for 2 min, then at 95°C for 15 min, followed by 45 cycles of denaturation at 95°C for 15 sec and annealing/elongation at 60°C for 30 sec. All reactions were carried out in duplicate per sample using sjTREC primers and sjTREC probe as well as albumin as a single-copy albumin gene to normalise the results, taking into account the amount of input DNA. During PCR, the amplification mediated the cleavage of the probes, which contain a quencher (TAMRA) and a reporter (FAM) dye. This, in turn, leads to the separation of the quencher from the reporter, thereby inducing fluorescence of the reporter dye. The expression of target (sjTREC and albumin) in analysed samples was established by measuring the threshold cycle (CT), defined as the cycle number at which the fluorescence generated by cleavage of the probe passes a fixed threshold above baseline. We calculated the standard deviation between the duplicates, and the results were accepted for further analysis when the standard deviation was ≤ 1. Next, we considered the average for all the duplicates with standard deviation ≤ 1, and the normalised sjTREC expression in each sample was calculated as a difference between average CT values of albumin and average sjTREC CT values.

### 4. Definition of telomere fast and slow agers and comparison between biological ageing methods

Telomere lengths were used to identify participants with above or below average telomere length. To do so, we fitted a linear model to each cell type’s telomere length using age, including months, as regressor. We then identified participants under one standard deviation of their predicted value. We considered all participants with at least two cell types passing the 1 standard deviation threshold to be fast or slow agers. Participants identified with some cell types above 1 standard deviation and some below were excluded. Thus, we defined two groups of participants: fast and the slow agers. We then performed a logistic regression using either sjTREC or methylation age as a regressor explaining this binary category (fast agers or slow agers) while controlling for age, sex and cell counts.

### 5. GREML heritability estimation

We used the GCTA^40^ software for narrow sense heritability estimation. We used the microarray SNP data available in LLD to calculate a genetic relationship matrix (GRM) using variants with a MAF > 1% (GCTA --make-grm). Next, we used the estimated GRM to calculate the amount of variance explained by the random effect of genetics in a linear mixed model (GCTA --reml) while accounting for sex and age as fixed-effect covariates.

### 6. GWAS of telomere length

Using a previously described pipeline^103^, we performed an association analysis of telomere length of each cell type with genome-wide SNP genotype data by calculating the Spearman correlation coefficient between telomere length measurements corrected for age and sex and SNP genotype dosages. We corrected the results for multiple testing by permuting genotype labels 10 times to create a null distribution that was used to control FDR at 0.05.

### 7. Polygenic risk score calculation

The contribution of genetics was calculated using a PRS created using 20 independent genome-wide significant loci reported a large European-descent telomere GWAS study^15^. A weighted PRS was calculated for each LLD participant using PLINK 1.9 --score sum function^104^. The PRS was used as a regressor using chronological age as covariate and telomere length of each cell type as dependent variables of six different linear models.

### 8. Phenotype correlations

We preselected 90 phenotypes for correlation with telomere length based on the relevance and sample size of each phenotype. We used telomere length as the dependent variable while the phenotype was standardised and used as the explanatory variable. Linear models were fitted in R (v4.0.1) by ordinary least squares (OLS). We fitted all models while accounting for the effect of age and sex. We then fitted a second model that also accounted for the effect of different blood cell counts available per sample, namely granulocytes (basophils, eosinophils and neutrophils), erythrocytes, lymphocytes, monocytes and thrombocytes measured per plasma litre. We computed FDR from all P-values using the Benjamini-Hochberg procedure^105^ as implemented in R base functions.

For some specific diabetes-related and BMI-related phenotypes, such as “Waist circumference in cm” and “Body Mass Index (kg/M^2^)”, we included paternal and maternal age and parental smoking habits as additional covariates in the model discussed above. At the same time, to disentangle maternal and paternal age effects, we applied a linear model including covariates (age, sex, cell counts) and both phenotypes, using each telomere length as dependent variable. We included an l1 penalisation (Ridge regression) using glmnet in R^106^ to shrink the estimates of superfluous covariates. The strength of the regularisation parameter (λ) was estimated by a 10-fold cross-validation.

To identify possible confounders in BMI-related phenotypes, we performed correlations between BMI and parental smoking and parental age using an additional 10,000 participants from the Lifelines study population^41^.

### 9. Assessment of total variance explained by phenotypes

To assess the total variability explained by each layer of information, we selected all phenotypes found significantly associated with at least two telomere lengths under an FDR of 0.05. We then fitted five nested models. First, we regressed out the effect of chronological age in telomere length. Next, we fitted three nested models with intrinsic host factors: one adding sex, a second adding cell counts and a third adding waist circumference and BMI. These were fitted using a lasso shrinkage model (using glmnet in R^106^) to explain the residuals of telomere length after removing the chronological age effect. The fourth model added the parental phenotypes age and smoking to the intrinsic factors model. The final model added the PRS (see *Polygenic Risk Score calculation*) on top of the parental model. We computed the variance explained (R^2^) by each model and the ΔR^2^ of the variability gained by each layer of information introduced.

### 10. Methylation association analyses

We performed a GWAS of all probes with parental phenotypes using OLS in R. The following five phenotypes were used: age of mother and father at participant’s birth, maternal smoking during childhood, maternal smoking during pregnancy and paternal smoking during childhood, all while correcting for age, sex and cell counts as covariates. Bonferroni multiple testing correction was applied to correct for 5x418,499 = 2,092,495 tests. Next, we assessed the association of methylation and telomere length using a similar linear model, adjusting for age, sex and cell counts as covariates.

### 11. Mediation analysis

To test if the parental phenotype effect on telomere length may be mediated by methylation of certain probes, we performed a mediation analysis. For this we selected triplets of parental phenotypes (independent variable), methylation probes (mediator) and telomere length (dependent variable) using the following criteria: We required that a parental phenotype was associated with the methylation probe with FDR < 0.05, that methylation at this probe was associated with telomere length at a nominal p-value threshold of 0.05 and that the parental phenotype was associated with telomere length at a nominal p-value threshold of 0.05. Next, we estimated mediation using the mediation R package^107^, adding age, sex and cell counts as covariates to all models. We performed a Benjamini-Hochberg multiple testing correction on the average causal mediation effect (ACME) p-value and required that the FDR < 0.05 and the average direct effect (ADE) p-value was > 0.05 (meaning that the parental phenotype effect is mostly mediated by methylation). In addition, we investigated a slightly different situation where methylation and telomere length are swapped. Here we checked if the parental phenotype effect on methylation is mediated by telomere length. While the real causality directions are very complex and often contain loops, and as a consequence these swapped ACME P-values are often < 0.05, we required that a swapped ACME p-value be larger than the main original ACME p-value and that the mediated proportion of the effect for the original direction be larger than for the swapped scenario.

### 12. Differential expression analysis at the single-cell level

#### scRNA-seq dataset features: library preparation, sequencing, alignment, pre-processing and quality control

To study gene expression changes with telomere length at the single-cell level, we used a subset of previously processed scRNA-seq data^49^ on unstimulated PBMCs from 62 LLD donors for whom we collected Flow-FISH telomere length data for at least one cell type in the current study. This scRNA-seq data was generated 5 years after collection of the Flow-FISH telomere length data. In short, scRNA-seq data was generated using the 10X Chromium Single Cell 3’ V2 chemistry and libraries were 150bp PE sequenced on the NovaSeq6000. The Cellranger v3.0.2 pipeline was used with default parameters to demultiplex, generate FASTQ files, map reads to the GRCh37 reference genome and generate a unique molecular identifier (UMI)-corrected count matrix per cell^108^. After quality control, 54,373 cells remained. In the original dataset, k-nearest neighbour clustering was used to cluster the cells.

We then performed automated cell type classification using Azimuth to annotate the cells^50^. In detail, we conducted a supervised analysis guided by a reference dataset to enumerate cell types that would be challenging to define with an unsupervised approach. Thus, we mapped our scRNA-seq query dataset to a recently published CITE-seq reference dataset of 162,000 PBMC measured with 228 antibodies^50^. For this process, we used a set of specific functions from the Seurat R package v4.0.0^50, 109^. First, we normalised the reference dataset using the SCTransform function. Then, we found anchors between reference and query using a precomputed supervised PCA transformation through the FindTransferAnchors function. Afterwards, we transferred cell type labels and protein data from the reference to the query. We also projected the query data onto the UMAP structure of the reference. For these two last steps, we used the FindTransferAnchors function. Finally, the high resolution cell-type-annotations predicted by Azimuth (celltype.l2) were combined in such a way to more closely reflect the resolution of the Flow-FISH annotations (i.e. naïve and memory CD4T and CD8T cells, NK- and B-cells) (**Supplementary Table 6**).

#### Differential expression analysis at the single-cell level

Telomere length DE analyses were conducted using MAST v1.16.0^110^ at single-cell resolution (sc-DEA) by selecting the matched telomere length measurement and gene expression level for each of these cell types. As a first approach (I), we performed an independent telomere length sc-DEA for each of the previously defined cell types (i.e. naïve and memory CD4T and CD8T cells, NK- and B-cells). We then conducted a second approach (II) in we combined multiple (sub)cell types together in the same analysis (i.e. we performed five different sc-DEA combining: all CD4T, CD8T, naïve T, or memory T and T-cells) while controlling for cell type annotation (**Supplementary Table 6**). As our first approach had indicated that we had insufficient cells to detect telomere length–associated DE genes in B- and NK-cells, we did not re-analyse these cells using our second approach. This strategy allowed us to increase the number of cells per donor per analysis and thereby increase the power to detect potential effects. In both cases, as input, we used the log-normalised and scaled expression counts from those genes expressed in at least 10% of the cells. MAST uses a two-part generalised linear model, specifically a Hurdle model, on zero-inflated continuous data in which the zero process is modelled as a logistic regression and the continuous process is modelled as a linear regression. To accommodate the complex experimental design while controlling for covariates, including both biological variables (sex, age, donor and cell type) and technical factors (cellular detection rate (CDR) and 10x lane/experimental batch), we fitted a general linear mixed model (glmer) of the form:

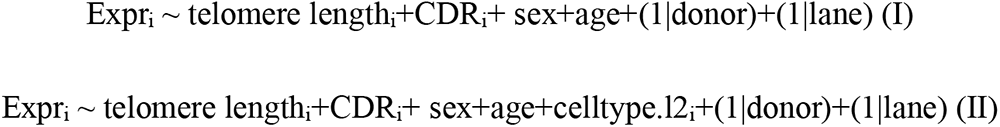

where Expr_i_ is the log-normalised, scaled expression of the gene being tested in cell i; telomere lengthi is the telomere length measurement using Flow-FISH in the tested cell type; CDRi is the fraction of genes that are detectably expressed in each cell; celltype.l2_i_ is the high resolution cell type annotation predicted by Azimuth; sex and age are phenotypic variables from the donors; donor is the donor LLD id and lane is the experimental batch, here being the lane of a 10X chip. First, we used the zlm function to fit a glmer (method = ’glmer’, ebayes=FALSE, fitArgsD=list, nAGQ=0, all other parameters set to default). Some of the gene models failed to converge when considering random effects (i.e. donor and lane), which resulted in NA estimated coefficients. To mitigate this convergence issue, nAGQ=0 was passed to the fitting function. We then used the summary function to perform a likelihood ratio test (LRT) to test for differences when we drop the telomere length factor (doLRT=telomere length, fitArgsD=list, nAGQ=0, all other parameters set to default).

Finally, genes were considered to be DE with telomere length when gene expression change was significant at an FDR < 0.05. The EnhancedVolcano function from EnhancedVolcano R package v1.8.0 was used to visualise the gene’s significance and log2FC. The pheatmap function from pheatmap R package v1.0.12 was used to visualise the expression pattern (log-normalised counts), clustering and annotation of the set of 97 unique DE among T-cells. For visualisation purposes, we down-sampled each of the Azimuth’s predicted T (sub) cell types to 100. To distinguish the four main expression patterns, the argument cutree_rows = 4 was set when using the pheatmap function. The pheatmap function was also used to visualise the LFC per telomere length unit among the set of 97 unique DE genes identified in T, CD4T and CD8T cells.

#### Literature analysis

To validate our sc-DEA findings, we compared our results to four biologically related studies that: i) revealed novel telomere proteins using *in vivo* cross-linking, tandem affinity purification and label-free quantitative LC-FTICR-MS^51^, ii) identified a set of genes associated with ageing in whole blood using bulk RNA-seq after correcting for cell type composition^53^, iii) created the GenAge human database as part of the HAGR databases composed of 307 human ageing–related genes^54^ and iv) defined an association between telomere length and methylated cytosine levels for both blood and EBV-transformed cell-line DNA samples^29^. To this end, we overlapped the set of 97 unique genes we found to be DE with telomere length against these four previously reported lists of genes and performed a two-sided Fisher’s exact test to assess whether these overlaps were significant. We could not test the overlap with ^29^ since multiple methylated CpG sites are linked to the same gene. For the tested overlaps, as our background list of genes, we gather the union of all tested genes in T-, CD4T and CD8T cells. As each study’s background list of genes, we considered all the tested genes in ^53^ and all the protein-coding genes (using Gencode v26, GRCh38 annotation) in ^51^ and ^54^.

#### Functional enrichment analysis

We performed a functional enrichment analysis through an over-representation analysis using WebGestalt (WEB-based GEne SeT AnaLysis Toolkit)^111^. As the input gene list, we used the 44 and 91 DE genes we identified in CD4T and all T-cells, respectively, split by their direction of effect (i.e. positively or negatively associated with telomere length). As the background gene set, we used the 2,426 and 2,427 expressed genes that were tested in the sc-DEA of the CD4T or all T-cells, respectively. As functional databases, we used two different pathway databases: KEGG and Reactome. Advanced default parameters were used (minimum and maximum number of genes for a category: 5 and 2000, multiple test adjustment: BH, significance level: FDR ≤ 0.05, number of categories expected from set cover: 10, redundancy reduction: affinity propagation and weighted set cover).

#### Differentially expressed genes telomere distance analysis

We compared the shortest distance to the telomere of DE genes compared to non-DE genes in CD4T cells and T-cells. Gene locations were retrieved using the human Ensembl BioMart dataset GRCh38.p13^112^. Chromosome length was used as a proxy for telomere location. We took the minimum between the start of the gene position and the distance between the end of the gene and the end of the chromosome as the distance from the gene to the telomere. We then performed a one-sided Fisher’s exact test to examine whether there was an enrichment of genes located at subtelomeric regions in the DE sets in comparison to background non-DE sets. We performed this analysis in positively and negatively telomere-associated genes separately and in each of the two cell types. To overcome a possible confounder effect of chromosome length, we subsampled with no replacement genes from the non-DE gene set while keeping the same chromosome proportions observed in the DE gene set, and again performed a Fisher’s exact test.

### 12. Survival analysis

We obtained survival information for 1,044 participants and ultimately the dates on which 11 participants had died. We measured the number of survival days since blood was donated. The R ‘survival’ package was used to perform a Cox regression analysis using survival (days) as dependent variable and telomere length as explanatory variable, while controlling for age, sex and blood cell counts. We tested the proportional hazards assumption, both of telomere length as a covariate and for the whole model (cox.zph function), using a p-value cut-off of 0.05.

## Supplementary Materials

**Supplementary Figure 1.**
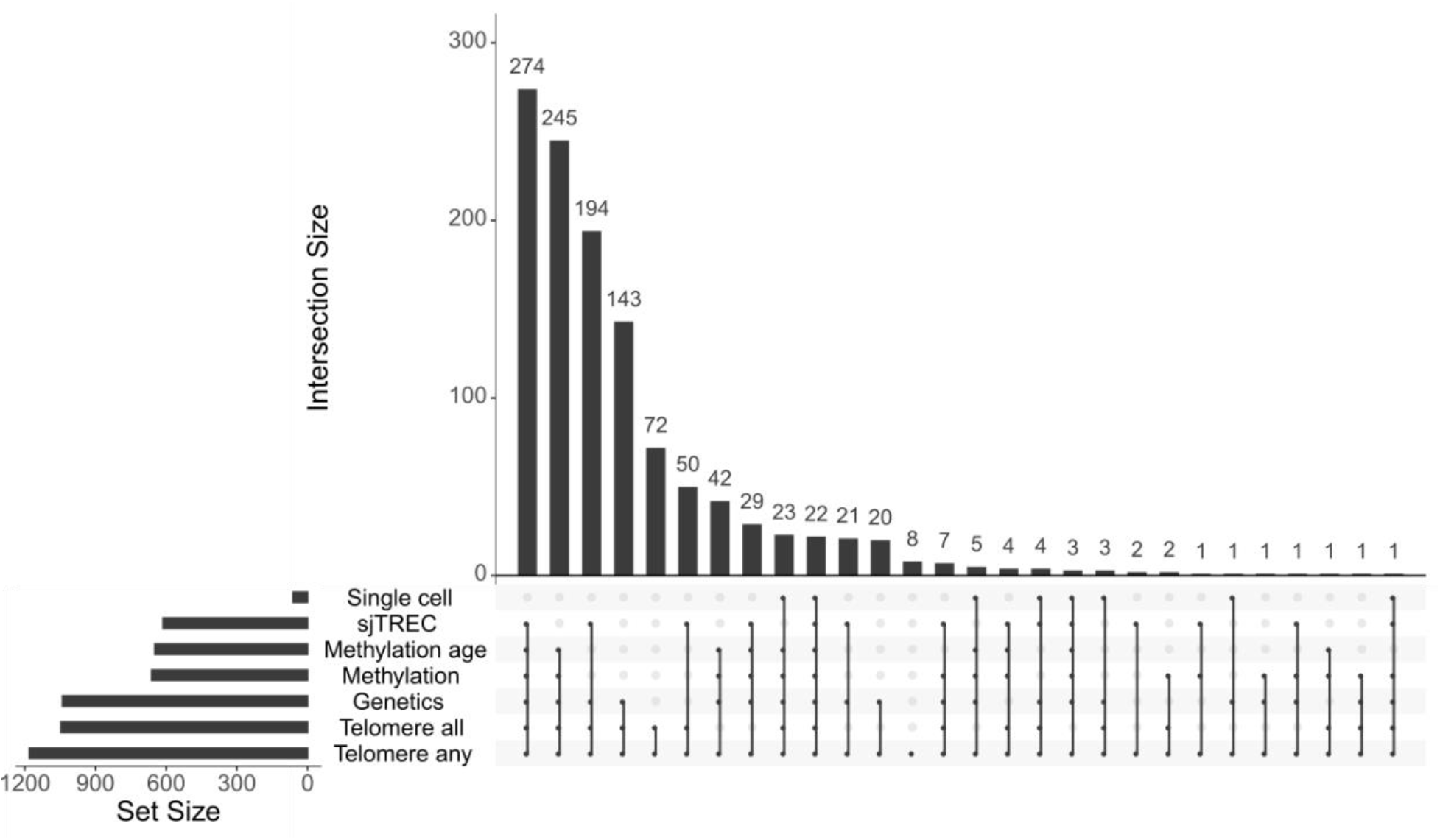
Phenotype markers in the LLD cohort. Representation of omics data used in this work. Horizontal bars indicate the number of Lifelines participants who had available information for the specific omics layer. Vertical lines indicate the number of samples in the intersection of the different layers and not seen in any other layer. Single cell (62) indicates the number of samples with single-cell RNA-seq data. sjTREC (613) indicates the number of people with qPCR measurements of sjTRECs. Methylation age (648) is a subset of participants where age was predicting using the methylation probes. Methylation (662) indicates the number of participants with available methylation arrays. Genetics (1,040) is the number of participants with available microarray data. Telomere any (1,180) are the number of participants with at least one cell population where telomere length was measured. Telomere all (1,046) are the number of participants with telomere lengths for all six cell populations available.

**Supplementary Figure 2.**
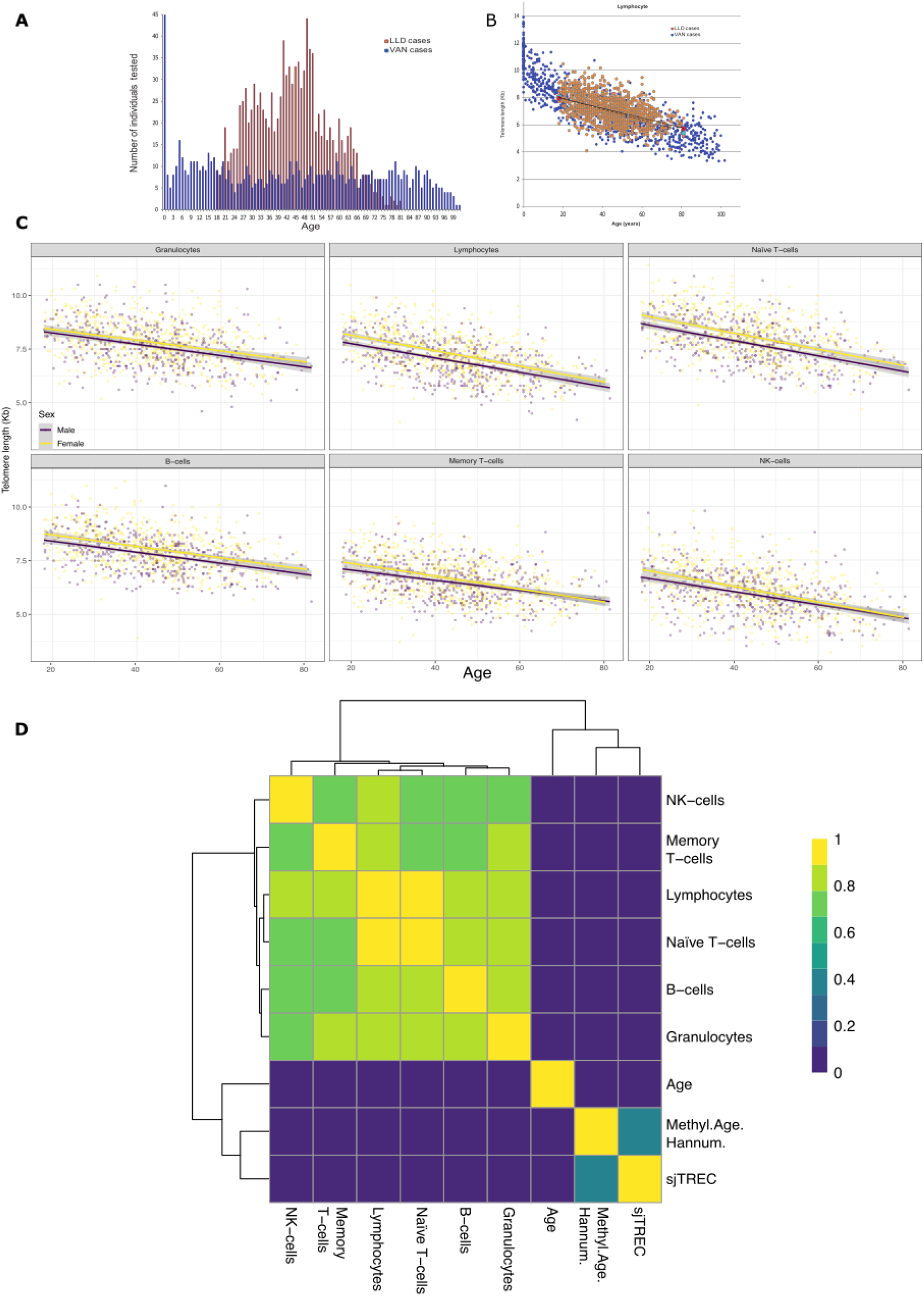
Sample cohort descriptions and measurements comparison. **A.** Number of study participants per age year in the Canadian study (VAN, blue, total n = 835, ages 18–81, n = 490) and the LLD study (LLD, orange, total n = 1049). **B.** Total lymphocyte cell type telomere length measurements (Kb) comparison between studies. Canadian cohort in blue. LLD cohort in orange. Each point represents an individual. 50^th^ percentiles of distribution are shown for each cohort and are consistent with one another (Canadian cohort light blue boundaries and dashed line, LLD cohort red boundaries and full line). The higher variability between models at older age is attributed to differences in study design and recruitment. **C.** Telomere length decreases with age in six different cell types. Each point represents a participant’s telomere length (Kb) in each cell type. Colour shows the sex of the participant. Trend line presents the average telomere length for a given age and sex group. Grey area surrounding trend lines indicates the 95% confidence interval of the trend line. **D.** Correlation (Spearman’s rho value) between the absolute value of telomere lengths, Hannum-based methylation age and sjTREC qPCR relative expression residuals after chronological age (Age) regression.

**Supplementary Figure 3.**
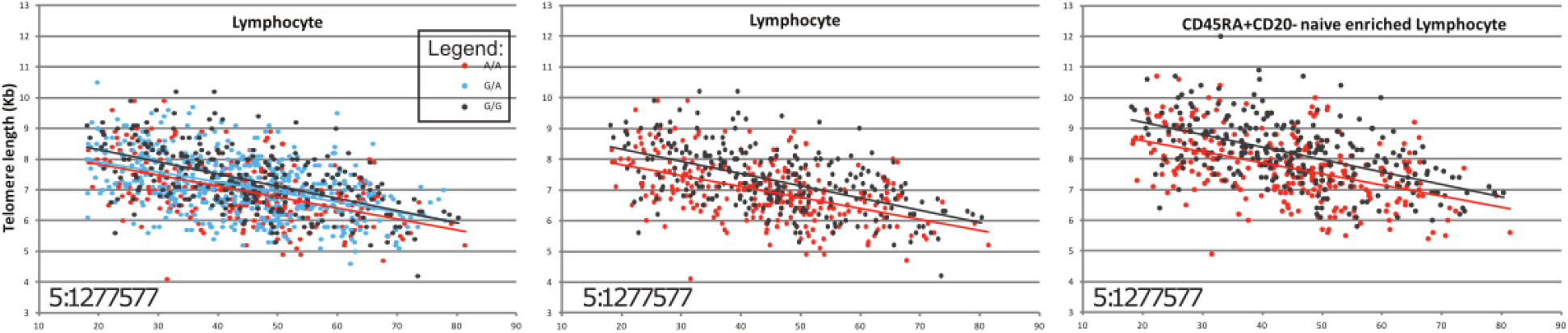
Telomerase SNP genotype with significantly different telomere length. *TERT* rs33961405 (chromosome 5:1277577) genotypes and respective linear regression are shown for the lymphocyte cell subset, with homozygous genotypes displayed for the lymphocyte and the CD45RA+CD20-naïve enriched cell type.

**Supplementary Figure 4.**
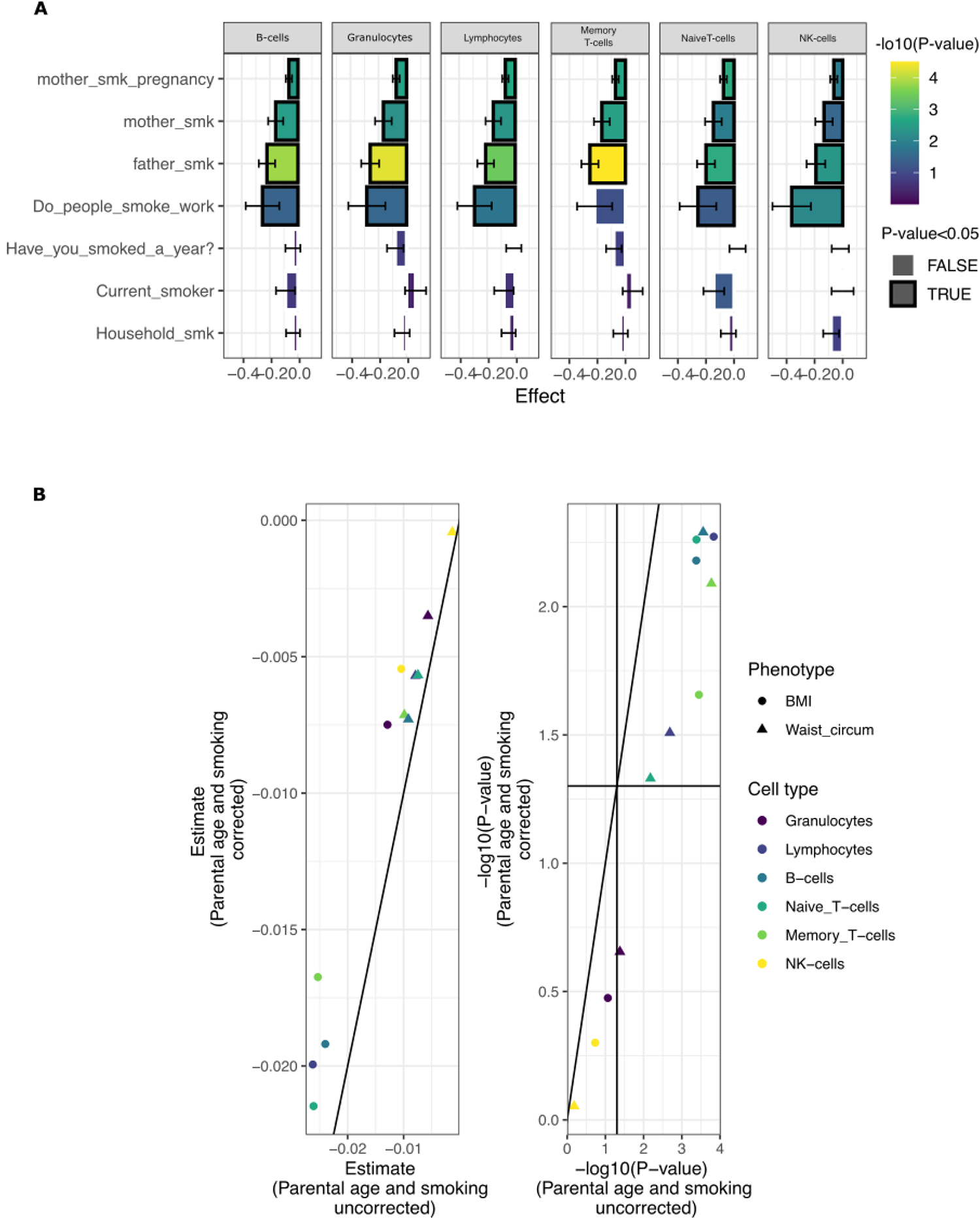
Phenotype exploration. **A.** Smoking effects on participant telomere length. Length of bar represents the linear model estimate. Colour shows the estimated P-value. Black outline indicates significant associations (nominal P<0.05). Error bars indicate the standard deviation of the estimation. Most phenotypes contain two levels (yes/no), except household_smk, which is a continuous variable of the number of people smoking at the participant’s residence, and mother_smk_pregnancy, which is a 4-level factor that includes: mother did not smoke, mother smoked before pregnancy, mother quit/reduced smoking during pregnancy and mother kept smoking regularly during pregnancy. **B.** BMI phenotypes corrected for parental age and smoking status. Left panel shows estimated effect size of BMI (circles) and waist circumference (triangles) on telomere length, while controlling for parental age and smoking status (y-axis) and without correction (x-axis). Associated -log10(P-values) of the estimates are shown in the right panel. Vertical and horizontal lines represent the nominal P-value significance threshold (0.05).

**Supplementary Figure 5.**
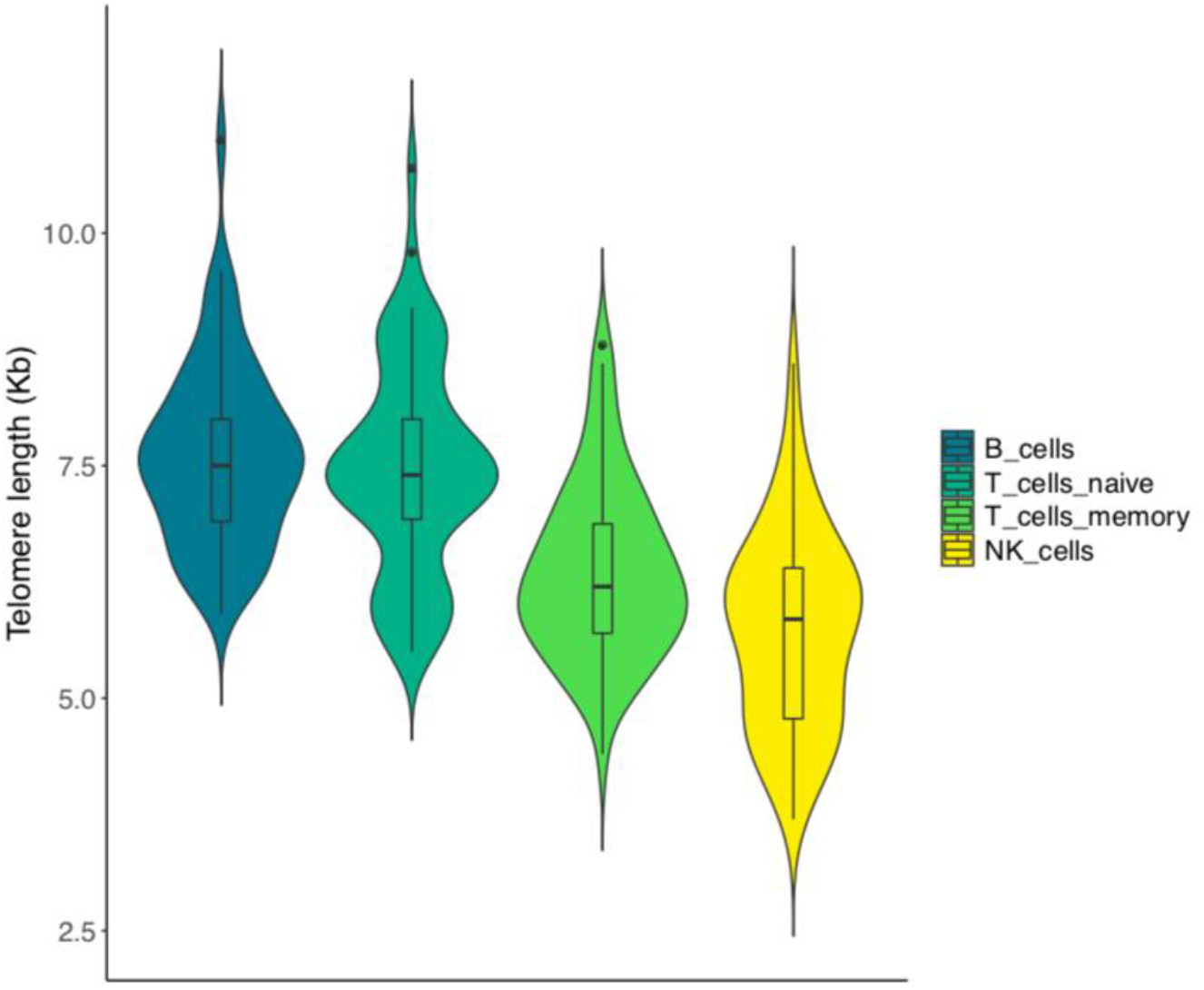
Telomere length in the subpopulations with single-cell expression data. Distribution of telomere length by cell type in the subset of 62 LLD donors for which both scRNA-seq and Flow-FISH telomere length data was collected. We confirmed that this subset of 62 LLD donors showed similar telomere length distributions to the total study population of 1,046 LLD donors in which telomere length was assessed (Figure 1A).

**Supplementary Figure 6.**
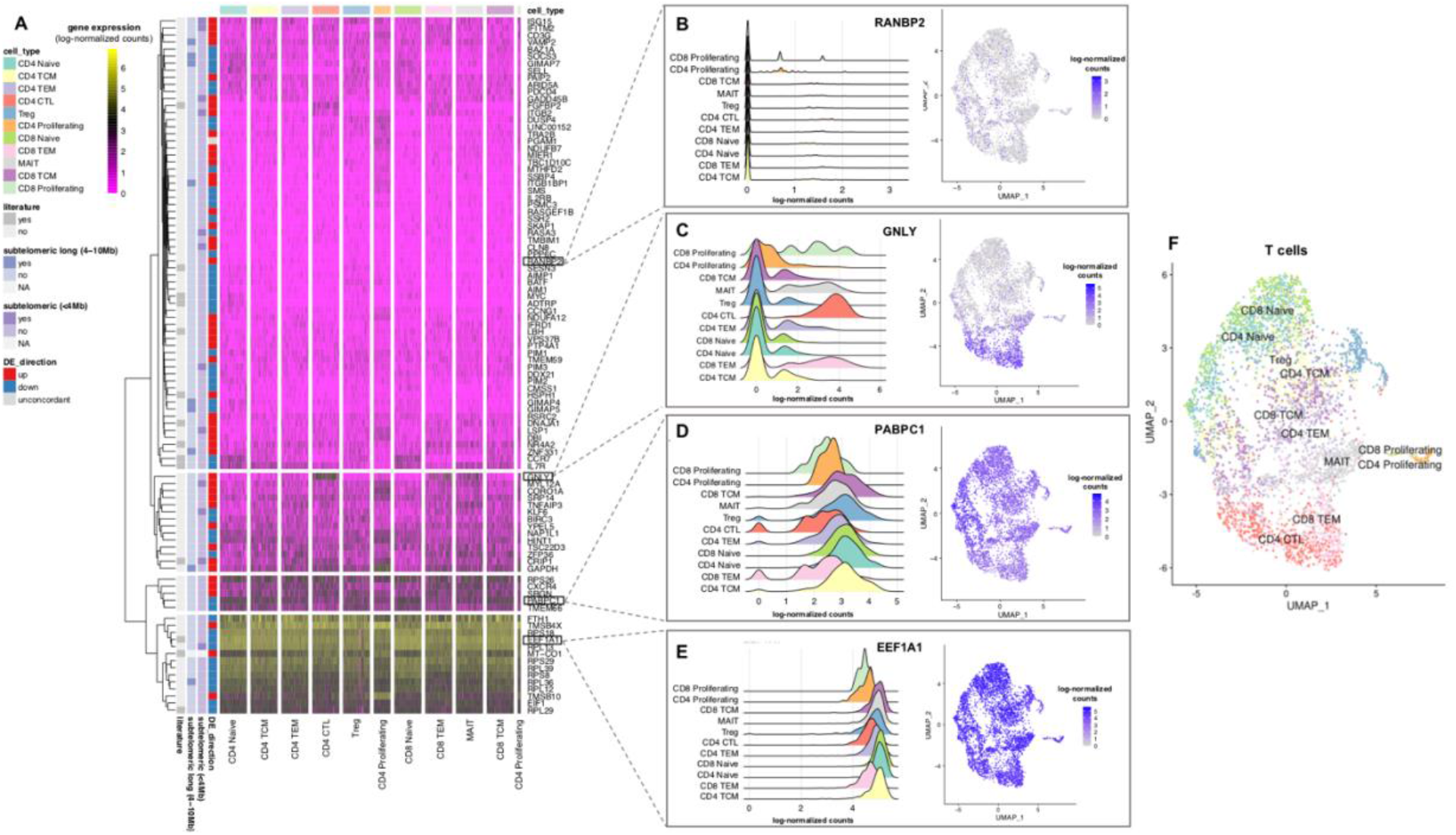
Gene expression pattern of the DEGs obtained through the DEA approach II across Azimuth’s predicted T-(sub)cell types. A. Heatmap of the gene expression (log-normalised counts) pattern of the set of 97 unique DEGs identified in T-, CD4T and CD8T cells. The hierarchical clustering of the genes revealed four main expression patterns. From bottom to top: genes highly and ubiquitously expressed, genes moderately and ubiquitously expressed, genes preferentially expressed in a particular cell type, and genes lowly and ubiquitously expressed. The column annotation bar shows all the cells from the same cell type. The row annotation bars show the DEG direction (DE_direction), whether the DEG is located at the subtelomeric (<4Mb) or subtelomeric long (4-10Mb) region, and if the DEG was previously reported in any of the following studies: Pellegrino-Coppola et al., 2021^53^, Tacutu R et al., 2018^54^, Buxton JL et al., 2014^29^ or Nittis T et al., 2010^51^ (literature) (**Supplementary Table 8**). Only one DEG (PGAM1) showed a different direction among T, CD4T and CD8T DEAs (DE_direction = unconcordant). The distance to the telomeres was not calculated for the mitochondrial gene MT-CO1 (subtelomeric (<4Mb) and subtelomeric long (4-10Mb) = NA). For visualisation purposes, we down-sampled each of the cell types to 100 cells. B-E. Examples of the four main gene expression patterns described in (A). Left: Gene expression (log-normalised counts) distribution plots across T (sub) cell types. Right: T-cells UMAP plots coloured by gene expression (log-normalised counts). F. T-cells UMAP plot coloured by Azimuth’s predicted T (sub) cell types. In B-F, we down-sampled each of the cell types to 500 cells for visualisation purposes.

**Supplementary Figure 7.**
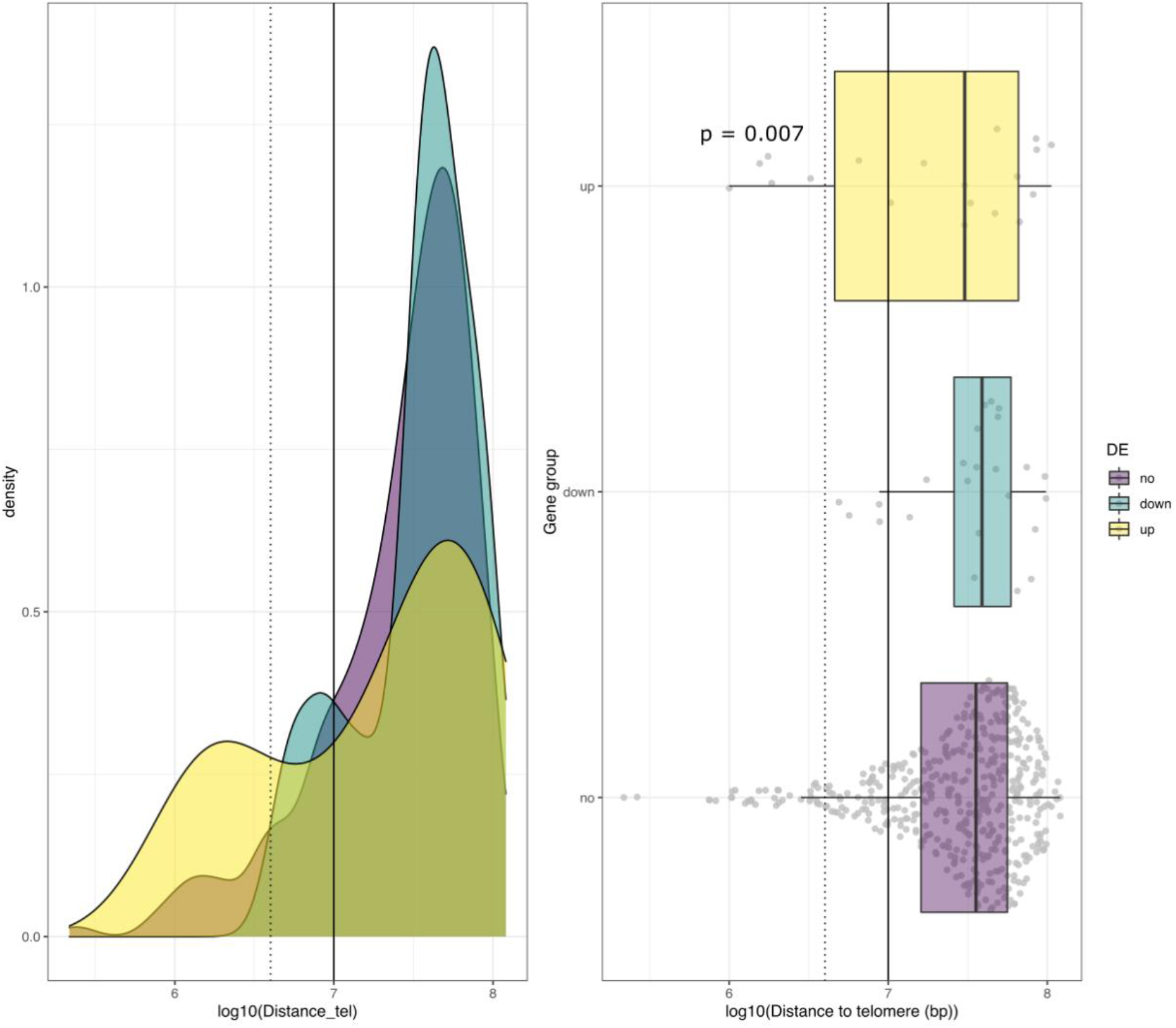
Distance to telomere of DE gene sets. Comparison of distance to the telomeres between DE and background genes in CD4 T-cells. A. Distribution of distance to telomere of DE genes (downregulated (blue) and upregulated (yellow)) and background (purple). B. Tukey boxplot of background genes, genes negatively associated with telomere length and genes positively associated with telomeres. Vertical lines represent subtelomeric region length (dotted, 4Mb) and maximum distance where TPE-OLD has been detected (not dotted, 10Mb).

**Supplementary Table 1. Descriptive statistics (mean age and BMI and female proportion) and number of participants in each of the data layers used in the study.**

**Supplementary Table 2. Telomere genome association summary statistics. Table 1.1** Replication of SNPs reported in Li et al.^15^. **Table 1.2** Summary statistics of associations which p<1×10^-5^. *EA:* effect allele, *beta:* estimated effect of effect allele load in telomere length, *SE:* standard error of the estimated effect.

**Supplementary Table 3. Exploratory statistics of phenotypes used.** *NA*: number of missing values. *Not NA*: number of samples used for analysis of the phenotype. *Info:* Maximum, minimum, mean and standard deviation of continuous phenotypes. For categorical phenotypes, it counts the number of participants per category.

**Supplementary Table 4. Summary statistics of associations between telomere length and phenotypes. Table 4.1** Summary statistics of the model controlling for sex and gender. **Table 4.2** Summary statistics of the model controlling for sex, gender and blood cell composition. *Regressor:* phenotype which effect the summary statistics makes reference to. *Dependent:* telomere length cell type used a dependent variable. *N:* number of samples used for assessment. *Levels:* number of levels (if categorical variable). *Level_n:* number of samples per level (if categorical) or number of samples (continuous). *FDR:* false discovery rate estimated using Benjamini-Hochberg.

**Supplementary Table 5. Methylation and mediation analysis summary statistics.** *Probe:* methylation site assessed; *eQTM_gene_p0.05:* gene which expression is associated with methylation of this probe with a p-value < 0.05^101^; mtl: telomere length measurement; {probe,pheno,mtl}-{probe,pheno,mtl}: association estimated effect between methylation (probe), phenotype (pheno) or telomere length(mtl); *fit.totaleffect*: association p-value between telomere length (outcome) and phenotype (predictor); *fit.mediator*: association p-value between methylation (outcome) and phenotype (predictor), *fit.dv*: association p-value between telomere length (outcome) and phenotype (predictor) when methylation is added as one of predictors; *pcor*: perarosn correlation, pheno = phenotype, mtl=telomere length, probe=methylation site; *ACME* (average causal effects): pval (pvalue), BH(Benjamini-Hochberg FDR), estimate (effect estimate); *ADE* (average direct effects); *Prop_med*: proportion of the effect of phenotype on telomere length that goes through methylation (mediator); *ACME_pval_reverse*: average causal mediation effects p-value for the opposite direction (effect of phenotype on methylation is mediated by telomere length); *ADE_pval_reverse*: average direct effects p-value for the opposite direction (effect of phenotype on methylation is mediated by telomere length); *Prop_med_reverse*: proportion of the effect of phenotype on methylation that goes through telomere length; *pass_filter*: Boolean value showing if the mediation result passes all filters

**Supplementary Table 6. Cell type classification.** Relationship between the cell type classification used for the two differential expression analysis approaches (DE_approach_I and DE_approach_II) and the automated high resolution (Azimuth_celltype_l2) cell type annotation using Azimuth. The number of cells (*n*) within each of the Azimuth_celltype_I2 cell type–annotations is shown. The Azimuth_celltype_I2 cell types are sorted from high to low *n*, within each of the levels of the DE_approach_I cell type resolution.

**Supplementary Table 7. Differential Expression Analysis (DEA) results.** Summarised model features from a ZlmFit object using MAST. **Table 7.1** DEA results for DE method 1. **Table 7.2** DEA results for DE method 2. *Primerid*: the gene; *coef*: point estimate; *ci.hi* and *ci.low*: upper and lower bound of confidence interval, respectively; *z:* z score (coefficient divided by standard error of coefficient); *Pr(>Chisq)*: likelihood ratio test p-value*; fdr*: False Discovery Rate, multiple testing correction method. Columns contain NAs if they are not applicable for a particular component or contrast.

**Supplementary Table 8. Differentially Expressed Genes (DEGs) features.** Summary information table for the unique DEGs (*unique_DEGs*) in each of the cell types (*group*) discovered using the DE_approach_II. For each unique DEG, we report information on their DE direction (*DE_direction*), whether the DEG is located at the subtelomeric (<4Mb) or subtelomeric long (4-10Mb) region and if the DEG was previously reported in any of the following studies: Coppola et al., 2021, Tacutu R et al., 2018, Buxton JL et al., 2014 or Nittis T et al., 2010 (*literature*).

**Supplementary Table 9. Functional Enrichment Analysis (FEA) results.** FEA results obtained through an over-representation analysis (ORA) by WebGestalt (WEB-based GEne SeT AnaLysis Toolkit) using the set of positively or negatively associated (as specified in the DE_diretion column) DEGs with telomere length in CD4T and T-cells (as specified in the cell_type column) by DEA approach II.

